# Tmem127-mediated immune receptor degradation regulates T cell homeostasis through the common gamma chain

**DOI:** 10.1101/2025.07.01.662281

**Authors:** Marko Hasiuk, Arlinda Negraschus, Denis Seyres, Romina Marone, Gytis Jankevicius, Lena Siewert, Christoph Schultheiss, Aida Muñoz Blázquez, Mascha Binder, Anne-Katrin Pröbstel, Sebastian Hiller, Vigo Heissmeyer, Lukas T. Jeker

## Abstract

Maintenance of T cell population size, which is important for immune homeostasis, is controlled by interleukin-7 (IL-7) and low-affinity TCR/MHC interactions that provide limited survival cues. Using arrayed CRISPR screening of miR-17∼92 targets, Bio-ID proximity labeling and proteomics we identified Tmem127 as an essential regulator of the T cell surface proteome. We validated interaction with the common gamma chain (IL-2Rγ) in a multi-protein complex. Tmem127 reduces IL-7 receptor surface expression to restrict homeostatic proliferation, thereby controlling naïve and central memory T cell population sizes. Tmem127 germline knockout (KO) mice display splenomegaly, accelerated experimental autoimmune encephalomyelitis and Tmem127-deficient bone marrow displays a competitive advantage over wildtype cells. Thus, we identified Tmem127 as an important regulator of the common gamma chain and immune homeostasis.

## Main Text

T cell homeostasis is achieved by a balanced system of signals that allows the maintenance of a stable and diverse population of T cells. Mice rely on thymic output to maintain their T cell numbers, while in humans, homeostatic proliferation replenishes the majority of the naïve T cell pool (*1, 2*). Two main signals that induce homeostatic proliferation of T cells are interleukin-7 (IL-7) signaling and low-affinity T cell receptor (TCR) interactions with self-major histocompatibility complex (self-MHC) molecules (*3–5*). Due to the limiting role of IL-7 in T cell homeostasis, its heterodimeric receptor consisting of the IL-7 receptor alpha chain (IL-7Rα; CD127) and the common gamma chain (IL-2Rγ; CD132; aka γc) is regulated on multiple levels. In a negative feedback loop, IL-7 signaling itself inhibits the expression of the Il7ra gene (*6*). In addition, IL-7 binding to its receptor causes rapid clathrin-dependent downregulation of the IL-7 receptor from the cell surface (*7*). In the absence of a signal, receptors undergo steady-state internalization at a slow rate, which balances the protein output, while signaling shifts the balance towards more rapid degradation (*8*). The γc is a shared subunit of the receptors for the whole IL-2 family of cytokines (IL-2, IL-4, IL-7, IL-9, IL-15, and IL-21) and is regulated by a constitutive clathrin-independent internalization (*9*). How the decision of recycling vs degradation of the receptor is made remains unknown. Upon binding of their specific cytokines, all IL-2 family cytokine receptors activate Janus kinases (JAKs), JAK1 and JAK3. JAK1 associates with the specific receptor subunits (e.g. IL7ra), whereas JAK3 associates with γc (*10, 11*). JAK1/3 phosphorylate the specific receptor subunits, which, through a cascade of events, results in phosphorylation, dimerization, and, ultimately, nuclear translocation of signal transducers and activators of transcription (STAT) proteins. IL-7 signaling activates STAT5, which directs downstream transcriptional responses to the cytokine signal (*12, 13*). Due to γc’s centrality to mediate multiple cytokine signals that affect innate and adaptive immune cells, including CD4^+^ and CD8^+^ T cells, loss-of-function mutations of γc lead to X-linked severe combined immunodeficiency (X-SCID) in humans (*14*). As a consequence, a γc-specific antibody that blocks signaling of all γc cytokines, depletes NK and T cells in non-human primates and improves immune-mediated disorders in mouse models of graft-versus-host disease (GVHD) and experimental autoimmune encephalitis (EAE) (*15*). Thus, γc is a key component of central cytokine receptors that regulate many immune cells and are involved in immune pathogenesis.

miR-17∼92 is a microRNA cluster known to regulate T cell homeostasis and function (*16*). Its overexpression in lymphocytes causes uncontrolled expansion of CD4^+^ T cells and fatal autoimmunity in mice (*17*). In fact, miR-17∼92 is also known as oncomiR-1 since it represents the first known oncogenic miRNA (*18*). Different miRNAs of the cluster regulate specific sets of genes and functions (*19, 20*). Conversely, miR-17∼92 deficiency causes B and T cell lymphopenia, and it regulates IL-7 receptor expression and IL-7 sensitivity by an unknown mechanism (*21, 22*). Thus, miR-17∼92 controls critical T cell functions by regulating genes that control cell cycle, survival, and T cell differentiation (*16*). However, beyond some well-validated target genes like Pten and Bim, for most target genes it remains largely unknown to what degree miR-17∼92-mediated regulation is functionally relevant and in addition, some phenotypic effects have not been ascribed to any target (*16*). We recently demonstrated that transgenic miR-17∼92 can functionally substitute for CD28-deficiency to a substantial degree (*23*). We identified 68 bona-fide miR-17∼92 target genes in T cells and hypothesized that deleting individual target genes in CD28KO CD4^+^ T cells should result in phenotypic rescue comparable to target gene repression by transgenic miR-17∼92. Indeed, CRISPR/Cas9-mediated deletion of individual miR-17∼92 target genes in naïve T cells differentially affected T cell proliferation and expression of T cell surface proteins (*23*). Here, we functionally assessed the miR-17∼92 target genes in naïve T cells using an arrayed CRISPR screen. We identified Tmem127, a transmembrane adaptor protein that was recently reported to participate in ubiquitination and lysosomal degradation of MHC (*24*), as an important regulator of key T cell surface proteins. The absence of Tmem127 leads to inappropriate γc expression and increased γc cytokine sensing which in turn results in hyperproliferation of naïve and central memory T cells. Tmem127 interacts in a multi-protein complex with γc to mediate its degradation. Thus, we uncovered a new mechanism by which Tmem127 limits γc surface protein abundance posttranslationally to actively restrain T cell homeostatic proliferation.

### Tmem127 restrains T cell surface proteins and proliferation of naïve CD4 T cells

To systematically explore the miR-17∼92 regulatory gene network in T cells, we screened all 68 target genes using an arrayed CRISPR/Cas9 loss-of-function screen in naïve, CD28KO CD4^+^ T cells. Aiming for a phenotypic rescue of CD28-deficiency, we used flow cytometry to quantify parameters that were restored by transgenic miR-17∼92 overexpression in CD28KO CD4^+^ T cells (*23*). The median guide RNA (gRNA) cutting efficiency measured by Sanger sequencing-based Tracking indels by decomposition (TIDE) was 26.25%, reaching 37.1% when only the best-performing gRNAs per gene were taken into account (fig. S1A) (*25*). The relatively low cutting efficiencies are likely attributed to the fact that we performed CRISPR/Cas9 KO in naïve T cells, before T cell activation. However, we recently demonstrated that even low KO frequencies are useful in this context since the CD28-deficiency converts the loss-of-function into a gain-of-function screen (*23*). gRNA performance did not correlate with predicted Rule set 2 scores (fig. S1B) (*26*). Principal component analysis (PCA) revealed that the majority of gRNAs clustered together, whereas a few distinct gRNA clusters were more separated. These included the non-targeting control (NTC) gRNAs in wildtype (wt) cells (“NTC_wt”) used as a positive control and, in the opposite direction on PC1, T cell receptor α KO (“TRACKO”) used as a negative control. A known miR-17 target, Jak1KO, clustered with TRACKO negative controls (Fig. 1A) (*27*). In contrast, CD45KO samples clustered closer to NTC_wt, reflecting CD45’s regulatory roles in TCR and cytokine signaling (Fig. 1A) (*28–30*). A distinct cluster separated by PC1 and PC2 was formed by PtenKO, possibly due to the previously observed hyperactivation and hyperproliferation. Finally, Nrbp1KO confirmed the recently reported upregulation of activation markers (Fig. 1A) (*23*). The most profound effect of a new miR-17∼92 target gene was achieved by Tmem127 targeting gRNAs. All 3 Tmem127 gRNAs clustered together with NTC wt cells during in vitro activation (Fig. 1A). Tmem127KO CD4^+^ T cells displayed increased surface expression of several immune proteins, affecting ICOS, CD25 and CD226 the most. These effects were specific since other parameters such as viability and PD-1 levels were unaltered (Fig. 1B). We confirmed the results in two independent screening experiments (Fig. 1C). Next, we analyzed the effect of Tmem127KO in naïve CD4^+^ T cells with intact CD28. Knockout of Tmem127 in T cells isolated from C57BL/6N mice neither affected the frequency of activated cells nor their activation-induced proliferation and led to only mildly increased expression of ICOS and CD44 during activation (fig. S1, C and D). Intriguingly, culturing naïve Tmem127KO CD4^+^ T cells with 5ng/ml IL-7 without in vitro activation resulted in significant upregulation of ICOS, CD25, and CD226 and a small but significant increase in the percentage of proliferating cells compared to non-targeted control cells (Fig. 1D). This suggests that T cell activation proceeds normally but steady state T cell phenotypes are altered by Tmem127 deficiency (fig. S1, C and D). Therefore, Tmem127 may play a more important role in naïve than activated T cells, as CRISPR-mediated Tmem127KO led to the upregulation of immune proteins on the surface of naïve CD4^+^ T cells and increased proliferation in vitro.

**Fig. 1.**
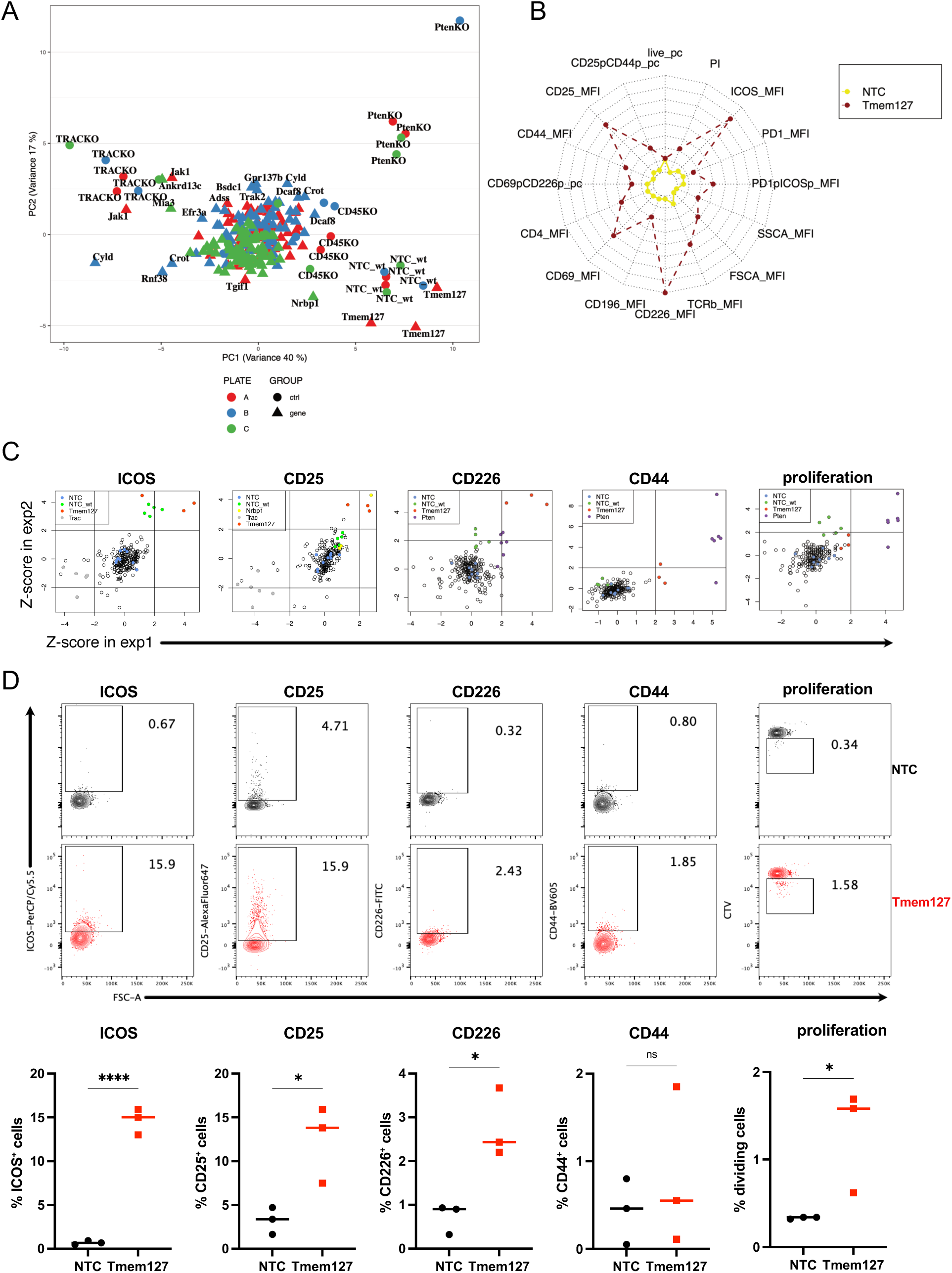
Tmem127 regulates surface levels of multiple T cell proteins. (**A**) Principal component analysis (PCA) of flow cytometry data from an arrayed CRISPR screening experiment, (**B**) Radar plot representing flow cytometry profiles of Tmem127KO (Tmem127) and a non-targeting control (NTC) sample from one CRISPR screening experiment, (**C**) Z-scores calculated from individual flow cytometry parameters (MFI, % of positive cells (pc) or proliferation index (PI)) representing two independent CRISPR screening experiments (exp1 + exp2), (**D**) Activation markers and proliferation of naïve CD4^+^ T cells after Tmem127KO or non-targeting control electroporation, line and dots represent the median and 3 independent experiments. P values were calculated using unpaired t-test. ns, not significant; *p < 0.05; **p < 0.01; ***p < 0.001; ****p<0.0001.

### Tmem127KO mice develop splenomegaly and display expansion of naïve T cells

To investigate Tmem127’s function in more detail, we generated germline knockout mouse strains. Oocytes were injected with spCas9-HiFi ribonucleoproteins (RNPs) targeting Tmem127 using the most efficient Tmem127 gRNA from our screening (Tmem127_1). The non-homologous end joining repair resulted in various alleles displaying deletions, insertions, or combinations thereof. We backcrossed 3 F0 mice containing 5 mutations to C57BL/6N mice. By intercrossing the F1 offspring, we established 5 distinct mouse strains carrying homozygous mutations (fig. S2A). Strains 1, 3, and 4 carried frameshifts and strain 2 displayed a substantial 214bp deletion in exon 2. In contrast, strain 5 displayed a 3bp in-frame deletion. Therefore, strains 1-4 were most likely disrupting Tmem127 function whereas the functional consequence for strain 5 remained unpredictable. Since ICOS was the most highly upregulated surface marker in miR-17∼92 transgenic CD28KO “rescue” mice (*23*) and ICOS was upregulated by CRISPR/Cas9-mediated Tmem127KO in vitro (Fig. 1D), we screened ICOS expression on T cells from all 5 newly generated mouse strains. Strains 1-4 displayed increased ICOS expression in naïve CD4^+^ and CD8^+^T cells compared to control littermates, whereas strain 5 didn’t display ICOS upregulation. Hence, the 3bp in-frame deletion is probably tolerable for the function of the protein (fig. S2B). We chose strain 1 (p.L56fs) carrying a 14bp frameshift deletion in exon 2 (c. 166_179del) for future experiments and will refer to it as “Tmem127KO” (fig. S2A). Tmem127KO mice are slightly smaller and weigh less, which is in line with a previously reported Tmem127 germline knockout strain (fig. S2C) (*31*). Despite the decreased overall body size and weight, Tmem127KO mice develop splenomegaly and increased spleen cellularity (Fig. 2A). In parallel, the frequency of αβT cells in the spleen of Tmem127KO mice was increased (Fig. 2B) with a decreased CD4/CD8 ratio (Fig. 2C). While there was no change in Tmem127KO naïve/memory cell frequencies, the absolute cell numbers of naïve CD4^+^, naïve CD8^+^ and memory CD8^+^ T cells were increased in Tmem127KO spleens (Fig. 2D). All populations that expanded in Tmem127KO mice express CD62L, which explains the shift in CD4/CD8 ratio – the biggest memory CD4^+^ population is CD62L^-^. Because a change in T cell numbers can be a result of increased proliferation or survival in the periphery or increased thymic output, we compared the thymus size and development of T cells. We did not find any significant differences except a slightly increased frequency of cells in the gate containing both DN1 thymocytes and stromal cells (fig. S2, D to F). Of note, the CD4/CD8 ratio was not altered in the thymus, suggesting that changes in the periphery accounted for the shifted splenic CD4/CD8 ratio. Thus, the T cell expansion observed in Tmem127KO animals affected mature T cells but had minimal effects on thymic T cell development.

**Fig. 2.**
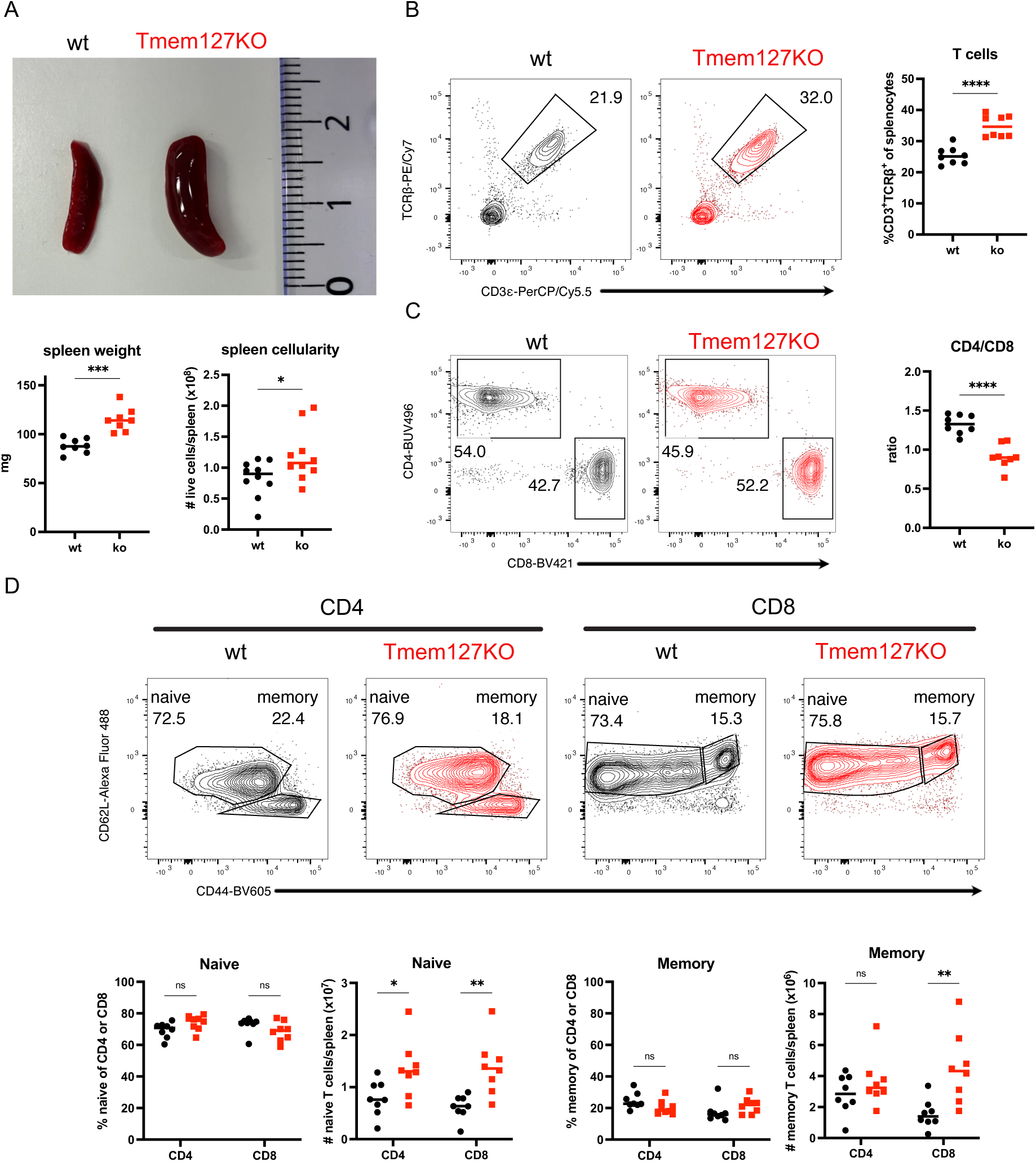
Phenotypic characterization of Tmem127KO mice. (**A**) Photo and quantification of weight and cellularity of spleens from Tmem127KO and wildtype (wt) mice, (**B**) Frequency of T cells among splenocytes from Tmem127KO and wt mice, (**C**) Ratio between CD4^+^ and CD8^+^ splenic T cells from Tmem127KO and wt mice, (**D**) Frequencies and cell counts of naïve and memory T cell populations from Tmem127KO and wt mice, dots represent mice and line median of 4 or 5 (A, spleen cellularity) independent experiments. P values were calculated using unpaired t-test or two-way ANOVA with Sidak’s multiple comparison test. ns, not significant; *p < 0.05; **p < 0.01; ***p < 0.001; ****p<0.0001.

### Tmem127 interacts with γc to mediate its posttranslational repression

After confirming Tmem127’s important function for immune homeostasis, we investigated the molecular mechanism. Tmem127 regulates surface protein abundance by promoting a protein’s ubiquitination and lysosomal degradation (*24, 32*). To systematically identify proteins regulated by Tmem127 in T cells, we compared the proteomes of wt and Tmem127KO from naïve and activated CD8^+^ T cells. In line with the results using CRISPR/Cas9 (Fig. 1D and fig. S1C) and the observation that naïve T cells were expanded in Tmem127KO mice (Fig. 2D), the effect of Tmem127-deficiency on the T cell proteome was more pronounced in naïve T cells than activated T cells (Fig. 3A and fig. S3A). A cluster of proteins that are normally expressed during activation were upregulated in naïve Tmem127KO CD8^+^ T cells (fig. S3A, red arrow). 202 proteins were significantly more abundant in naïve Tmem127KO CD8^+^ T cells compared to wt cells. An additional 116 proteins were only detected in Tmem127KO but not wt cells (Fig. 3B and fig. S3A, Suppl. Table 1). Among the 10 differentially expressed proteins with the most significant p values, nine were upregulated in Tmem127KO cells, consistent with a role for Tmem127 in protein degradation. Upregulation of H2-D1 confirmed the recent observation that Tmem127 is a major regulator of MHC expression (*24*). Furthermore, Rxrb (aka MHC class I Promoter binding protein), a nuclear receptor gene located in the MHC locus, was also upregulated. We validated MHC I (H2-D^b^, H2-K^b^) surface upregulation in freshly isolated naïve CD8^+^ Tmem127KO T cells by flow cytometry (fig. S3B). MHC class II (I-A) expression was increased when measured by flow cytometry, despite not being detected in the Bio-ID or proteomics data, possibly reflecting a higher sensitivity of flow cytometry (fig. S3B). Besides MHC, another noteworthy upregulated immune protein was γc (Fig. 3B). Next, we aimed to identify Tmem127 binding partners since it is an adaptor protein. We used Bio-ID proximity labeling to identify proteins in close proximity to Tmem127 (*33*). To discriminate T cell-specific interactions from more ubiquitous interactions we performed Bio-ID in CD8^+^ T cells and mouse embryonic fibroblast (MEF) cells. In addition, we used full-length Tmem127 and C-terminally truncated Tmem127 because Tmem127 interacts with the E3 ubiquitin ligase Wwp2 through its C-terminal end (*24, 32*). We identified many T cell-specific membrane-associated proteins, e.g. T cell receptor components (CD3γ, CD3δ, CD3ε, CD3ζ), T cell co-receptors (CD8α, CD8β), co-stimulatory and inhibitory molecules (CD2, CD27, CD28, CD86, Icam1, Icos, CTLA-4), LAT, CD44, Slamf1, integrins, MHC molecules (H2-K1, H2-D1, H2-T23) and cytokine receptor components (IL-2rb, IL-4R, Ifngr1, IL-27ra, γc, CD130/gp130), suggesting that Tmem127 is in close proximity to many important T cell surface proteins. We also noted several proteins associated with the endolysosomal system (Vamp2, Vamp3, Vamp5, Vps8, Vps45, Vps51) (fig. S3, C and D; Suppl. Table 2). Next, we overlaid proteins in proximity to Tmem127 that were also significantly upregulated in Tmem127KO in our proteomics dataset. Among the proteins identified in both cell types (CD8^+^ T cells and MEFs) were MHC I and Wwp2 (Fig. 3C). These two hits validated recent studies and thus served as positive controls (*24, 32*). In addition, we identified immune receptors such as Tgfbr2 and CD120b/Tnfrsf1b as well as multiple proteins related to the Golgi (Fam234b, Fam91a1, Zdhhc18, and Tmem87a) or endoplasmatic reticulum (Slc33a1) (Fig. 3C). We again used flow cytometry to validate expression of selected proteins. CD120b, CD130 and CD226 surface levels were increased in freshly isolated naïve CD8^+^ Tmem127KO T cells, confirming the quality of the proteomics and Bio-ID datasets (fig. S3E). Among the T cell-specific proteins were Icam1 and γc, which had one of the highest abundance ratios in the Bio-ID experiment (Fig. 3C). To investigate whether Tmem127 might directly bind γc, we used Alphafold 3 (AF3) modeling of protein interactions (*34*). We included Tmem127, γc, Susd6, and Wwp2 amino acid sequences in an effort to understand their potential interaction. The AF3 model predicted the assembly of a potential complex, with Tmem127 in the center, and Susd6 and γc on opposite sides of Tmem127 (Fig. 3D). Susd6 potentially provides an additional contact area with the extracellular part of the target. In summary, Tmem127-deficiency significantly altered the T cell proteome and proximity labeling detected Tmem127 in close proximity to important T cell surface proteins but also proteins of the endolysosomal system. An AF3 model predicts a multi-protein complex in which Tmem127 acts as a transmembrane adaptor directly interacting with γc as well as other proteins including Susd6 and Wwp2.

**Fig. 3.**
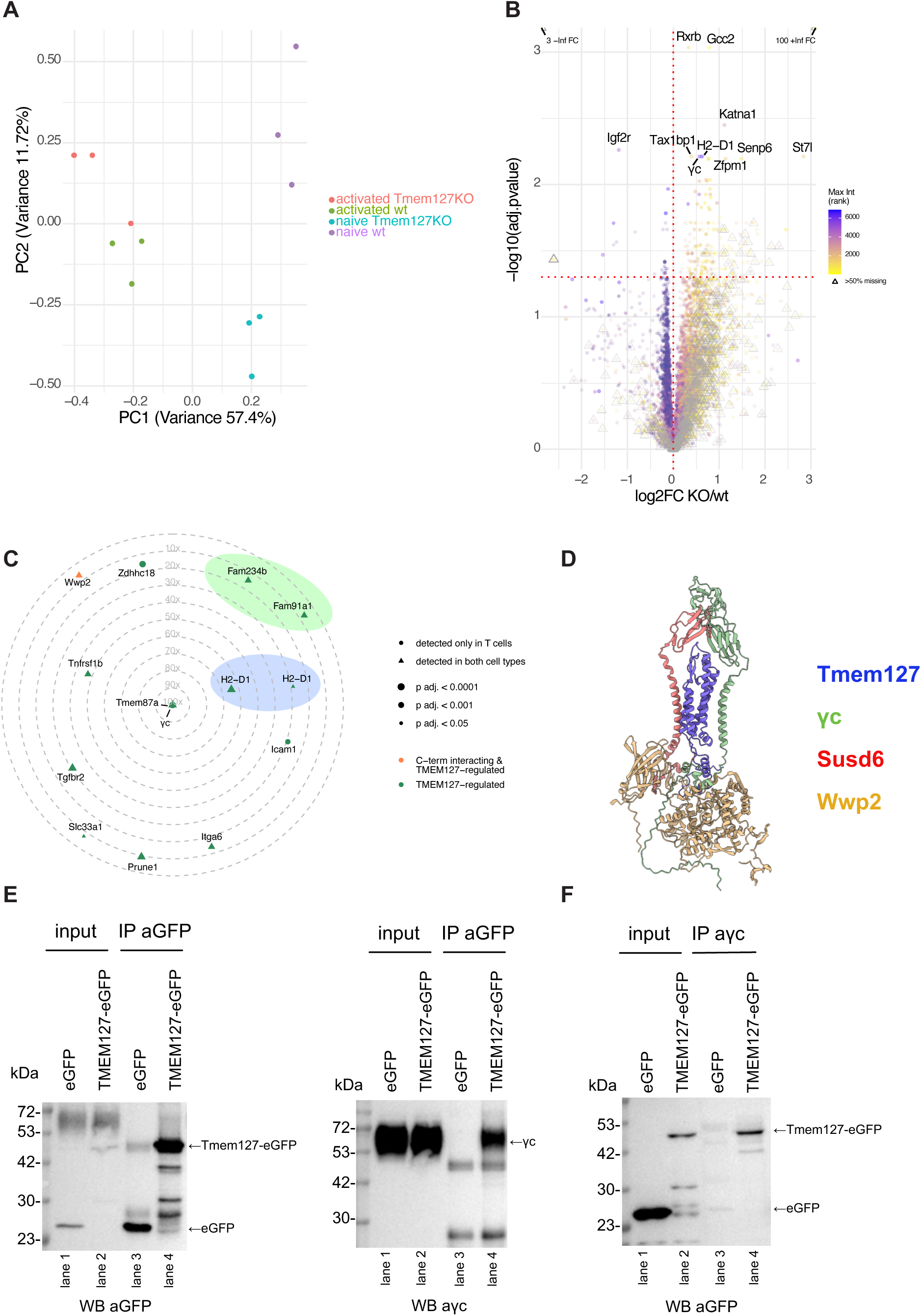
Identification of Tmem127 targetome using proximity labeling assay and proteomics. **(A)** Principal component analysis (PCA) of proteomics comparing naïve and in vitro activated CD8^+^ T cells from Tmem127KO or wt mice, **(B)** Volcano plot visualizing differentially abundant proteins detected by proteomics in naïve CD8^+^ T cells from Tmem127KO or wt mice, (**C**) Radar plot showing Bio-ID abundance ratios of proteins that were differentially abundant in proteomics, comparing detection in cells expressing full-length Tmem127-BirA versus the eGFP-BirA negative control. Included proteins met the following criteria: adj. p-value (Tmem127-eGFP vs. eGFP) < 0.05, abundance ratio (Tmem127-eGFP/eGFP) > 5.0, high FDR confidence, and > 1 unique peptide, with only proteins overlapping with proteomics considered, (**D**) Alphafold 3 model of Susd6/Tmem127/Wwp2 complex with γc, (**E**) Co-immunoprecipitation (Co-IP) of HEK293T cells co-overexpressing γc and Tmem127-eGFP or eGFP. GFP-tagged proteins were pulled down using anti-GFP, and Western blotting was first performed with anti-γc to detect interaction. An anti-GFP antibody from a different species than that used for the pulldown was then used to assess expression levels. Lane 1: input (eGFP), Lane 2: input (Tmem127-eGFP), Lane 3: capture (eGFP), Lane 4: capture (Tmem127-eGFP), (**F**) Co-immunoprecipitation (Co-IP) of CD8⁺ T cells overexpressing Tmem127-eGFP or eGFP. Endogenous IL2RG was pulled down using an anti-IL2RG antibody, and Western blotting was performed with an anti-GFP antibody to detect interaction. Lane 1: input (eGFP), Lane 2: input (Tmem127-eGFP), Lane 3: capture (eGFP), Lane 4: capture (Tmem127-eGFP).

We next investigated the role of the C-terminal Tmem127 end. Leveraging the Bio-ID data obtained with full-length Tmem127 and C-terminally truncated Tmem127, we concluded that most regulated proteins were detected in close proximity to Tmem127 in a C-terminus-independent way, except for Wwp2, which was biotinylated only if Tmem127 was expressed with its C-terminus (Suppl. Table 2). This is in line with our AF3 model and published data (Fig. 3, C and D) (*32*). In cancer cells, Wwp2 is recruited to Tmem127 to ubiquinate and degrade MHC (*24*). To test if Wwp2 is also functionally relevant in T cells, we validated in naïve CD4^+^ T cells that CRISPR/Cas9-mediated Wwp2 knockout phenocopied ICOS and CD25 upregulation as seen in Tmem127KO cells (Fig. 1, B and C), although to a lesser degree than observed in Tmem127KO (fig. S3F). Thus, some of the increased protein abundance observed in Tmem127KO T cells likely results from reduced ubiquitination (through Wwp2 or potentially other ubiquitin ligases) and lysosomal degradation of Tmem127-regulated proteins (*32*). This is further supported by the proteins associated with the endolysosomal system which we found in proximity to Tmem127 (fig. S3, C and D; Suppl. Table 2). To further test if lysosomal degradation also controls MHC expression in T cells, we cultured naïve CD8^+^ T cells in the presence of the lysosome inhibitor Bafilomycin A1. MHC I is one of the least stable T cell proteins with a very short half-life (*35, 36*), and indeed, we observed dose-dependent upregulation of MHC I on the cell surface upon Bafilomycin A1 treated T cells (fig. S3G). In addition, we observed increased Lamp1 staining intensity in naïve Tmem127KO CD8^+^ T cells compared to wildtype controls, reflecting increased endocytic vesicles in Tmem127KO cells (fig. S3H) and a blockage of their lysosomal degradation (*37*). Thus, Tmem127 regulates the T cell surface proteome by mediating active lysosomal degradation.

Finally, a major finding of our data was the identification of γc as a novel Tmem127-regulated protein. Its dysregulation (Fig. 3, B and C) could explain some of the phenotypic changes observed in Tmem127KO mice (Fig. 2). We therefore investigated whether γc interacts with Tmem127, guided by our Bio-ID data and Alphafold 3 modeling. We co-overexpressed Tmem127-eGFP + γc or eGFP + γc in HEK293 cells and then performed immunoprecipitation (IP) using anti-GFP antibodies. Subsequent staining with anti-GFP resulted in a distinct band at 27 kD in lane 1 and enrichment after IP in lane 3. Similarly, in the presence of Tmem127-eGFP, a signal was detected at a higher molecular weight band at 50 kDa (lane 2 and enrichment after IP in lane 4), reflecting the Tmem127-eGFP fusion size (Fig. 3E, left panel). The interaction of Tmem127 with γc was determined by detecting a strong signal in lane 4 and absence in lane 3 using anti-γc immunoblotting (Fig. 3E, right panel). γc expression was comparable in the input samples (lanes 1, 2) and the GFP signal in the IP samples (lanes 3, 4) was comparable as well. The additional bands at 25 kDa and 50 kDa most likely represent the heavy and light chains of the antibodies. Thus, this data strongly supported that Tmem127 interacts with γc. To test whether we could also detect an interaction in T cells with endogenous γc expression, we transduced CD8^+^ T cells with Tmem127-eGFP or eGFP. We then performed IP using anti-γc antibody-coupled beads. Western blots with anti-GFP showed a much lower signal in the input samples for the Tmem127-eGFP transduced cells than the eGFP transduced T cells (Fig. 3F, lanes 1, 2). Nevertheless, in the IP with γc pulldown, a GFP signal could only be detected in T cells transduced with Tmem127-eGFP but not eGFP alone (lanes 3, 4). Collectively, and together with the Bio-ID data and AF3 model, these experiments provide strong support for a direct Tmem127/γc interaction in CD8^+^ T cells. In summary, among the many newly identified Tmem127 interactors, we biochemically confirmed interaction of γc and Tmem127 (Fig. 3E, F).

### Tmem127-mediated γc downregulation restrains IL-7 signaling and proliferation in vitro

After having identified γc as a new Tmem127-regulated protein, we investigated the functional consequences of its deregulation in Tmem127KO T cells. The γc is a part of the IL-7 receptor and therefore its de-repression could explain the observed T cell expansion in Tmem127KO mice (Fig. 2). In accordance with the proteomics data (Fig. 3B, Suppl. Table 1), we validated by flow cytometry that naïve CD8^+^ T cells from Tmem127KO mice displayed increased expression of both IL-7 receptor components, IL-7Rα (CD127) and γc (CD132), respectively (Fig. 4A). In contrast, γc mRNA expression (RNA-Seq) was even slightly decreased in Tmem127KO, confirming posttranscriptional regulation (fig. S4A). IL-7 receptor hyperexpression by Tmem127KO cells resulted in increased phosphorylation of the downstream signal transducer Stat5 in response to IL-7 in vitro (Fig. 4B). This demonstrated that Tmem127KO CD8^+^ T cells were hypersensitive to IL-7. As a consequence, freshly isolated naïve CD8^+^ Tmem127KO T cells displayed a higher percentage of Ki-67^+^ cells (Fig. 4C), suggesting an increased homeostatic proliferation rate in vivo. To further test the in vitro proliferative capacity, we cultured naïve CD8^+^ Tmem127KO T cells with two IL-7 concentrations. Tmem127KO CD8^+^ T cells proliferated more than wt T cells when cultured with 5ng/ml but not 1ng/ml IL-7, measured by cell tracker violet (CTV) dilution (Fig. 4D) and Ki67 staining (Fig. 4E). Thus, we observed a dose-dependent propensity for hyperproliferation in response to IL-7 (i.e. an extrinsic signal) but at low IL-7 concentrations there was no cell-intrinsic hyperproliferation. Besides that, IL-7 signaling also led to the upregulation of Eomes and intracellular CTLA-4 in Tmem127KO cells (fig. S4, B and C). Tmem127KO cells also demonstrated pronounced transcriptome changes (fig. S4D). Tmem127KO led to decreased expression of multiple chemokines and chemokine receptors (Ccl3, Ccl4, Ccl5, Ccl6, Ccl9, Cxcl10, Ccr2, Ccrl2) and cytotoxicity-associated genes (Gzma, Gzmb, Klrk1, Klri2, Klrb1c, Klra4, Klra8, Klre, Klrb1b, Klra1) (fig. S4E). Reflecting their increased IL-7 sensitivity, Tmem127KO cells upregulated the expression of genes responsible for cell cycle progression and mitosis (S100a8, Nusap1, Cenpf) (fig. S4E). In addition, Tmem127KO cells exhibited upregulation of mRNAs that are known to be controlled by the transcription factor Myc (fig. S4F).

**Fig. 4.**
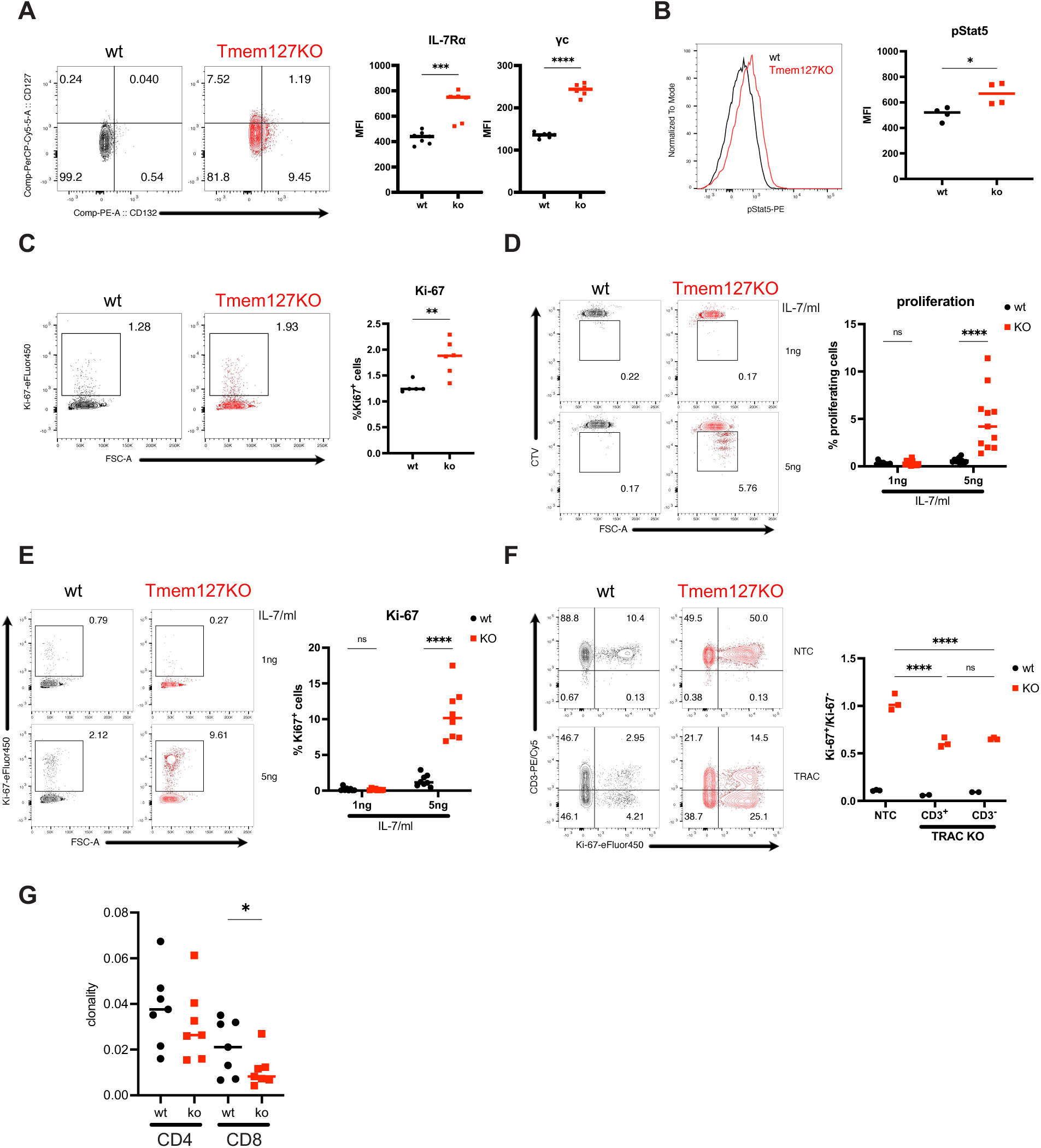
Tmem127KO cells are hypersensitive to IL-7 in vitro. **(A)**Surface level of IL-7 receptor on naïve CD8^+^ T cells from Tmem127KO and wt mice, **(B)** pStat5 staining after 1 day of in vitro IL-7 culture of naïve CD8^+^ T cells from Tmem127KO and wt mice, **(C)** Ki-67 levels in freshly isolated naïve CD8^+^ T cells from Tmem127KO and wt mice, **(D)** CTV dilution and, **(E)** Frequency of Ki-67-positive cells after 4 days of naïve CD8^+^ T cells in vitro culture with 1 or 5 ng/ml of IL-7 comparing Tmem127KO and wt mice, **(F)** Frequency of Ki-67-positive cells after 4 days of naïve CD8^+^ T cells in vitro culture with 1 or 5 ng/ml of IL-7, comparing TRACKO to non-targeting control electroporated Tmem127KO and wt cells, **(G)** Clonality of CD4^+^ and CD8^+^ T cells derived from TCRseq performed on splenocytes from Tmem127KO and wt mice, dots represent mice and line median of 2 independent experiments. P values were calculated using unpaired t-test or two-way ANOVA with Sidak’s or Tukey’s multiple comparison test. ns, not significant; *p < 0.05; **p < 0.01; ***p < 0.001; ****p<0.0001.

To discriminate whether the increased homeostatic proliferation of Tmem127KO cells was a result of their increased MHC surface levels or purely a result of increased IL-7 signaling, we used CRISPR/Cas9 RNPs to knock out the TCRα in wt and Tmem127KO T cells. As before, proliferation of Tmem127KO T cells was significantly higher than wt cells, both in T cells that received TCRα targeting CRISPR RNPs (“TRAC”) and controls (“NTC”). Increased proliferation of Tmem127KO T cells was identical in CD3^+^ and CD3^-^ cells (Fig. 4F). Thus, the increased proliferation was independent of TCR signaling and hence MHC expression. This confirmed that IL-7 signaling rather than increased MHC was a main driver of increased proliferation of Tmem127KO T cells. Intriguingly, both CD3^+^ and CD3^-^ populations proliferated less than unedited Tmem127KO cells, underlying the importance of TCR-MHC interaction for homeostatic proliferation (Fig. 4F). Finally, we hypothesized that the increased T cell proliferation could be driven by a few clones which would result in altered clonality. To test this, we performed TCR repertoire sequencing of naïve CD4^+^ and naïve CD8^+^ T cells isolated from spleens of wt and Tmem127KO mice. In line with the TRACKO results, we did not see increased clonality of the TCR repertoire that might arise from the TCR-mediated expansion of a few clones. Rather, we observed a slight decrease in clonality for CD8^+^ T cells but no statistically significant difference for CD4^+^ T cells (Fig. 4G). This data reflects the observed preferential expansion of CD8^+^ T cells and supports that T cell proliferation in Tmem127KO animals is polyclonal and driven by IL-7 but not TCR engagement. These results are in line with patients treated with IL-7 which resulted in proliferation of naïve T cells and a diversified TCR repertoire (*38*). In summary, the increase of γc surface levels in Tmem127KO T cells was caused by a posttranscriptional mechanism and was functionally relevant in response to IL-7 sensing but there was no cell-intrinsic hyperproliferation. Thus, IL-7 hypersensitivity rather than increased MHC levels in T cells was the main driver of proliferation in vitro and we observed indications of increased homeostatic proliferation in vivo.

### Tmem127 regulates autoimmunity and CD8^+^ T cell homeostatic proliferation in vivo

To further investigate Tmem127’s functional relevance in vivo, we induced EAE in Tmem127KO mice using MOG_35-55_, since γc blockade alleviates the severity of several mouse models of autoimmunity including EAE (*15*). Tmem127KO mice displayed a more rapid onset of clinical symptoms during EAE but clinical scores at peak disease were comparable to wt (Fig. 5A). This was paralleled by increased frequencies of activated IFNγ^+^CD44^+^ CD8^+^ T cells (Fig. 5B) in peripheral lymph nodes and spleens of Tmem127KO mice. Thus, the accelerated disease onset in mice with Tmem127-deficiency was opposite to the attenuated disease after γc blockade and might be explained by the increased starting naïve T cell pool and the increased γc expression. Th1 and Th17 cell frequencies were not changed in Tmem127KO (fig. S5, A and B). In line with the clinical data, we didn’t observe differences in T cell infiltration into CNS at peak disease on day 16 (fig. S5C).

**Fig. 5.**
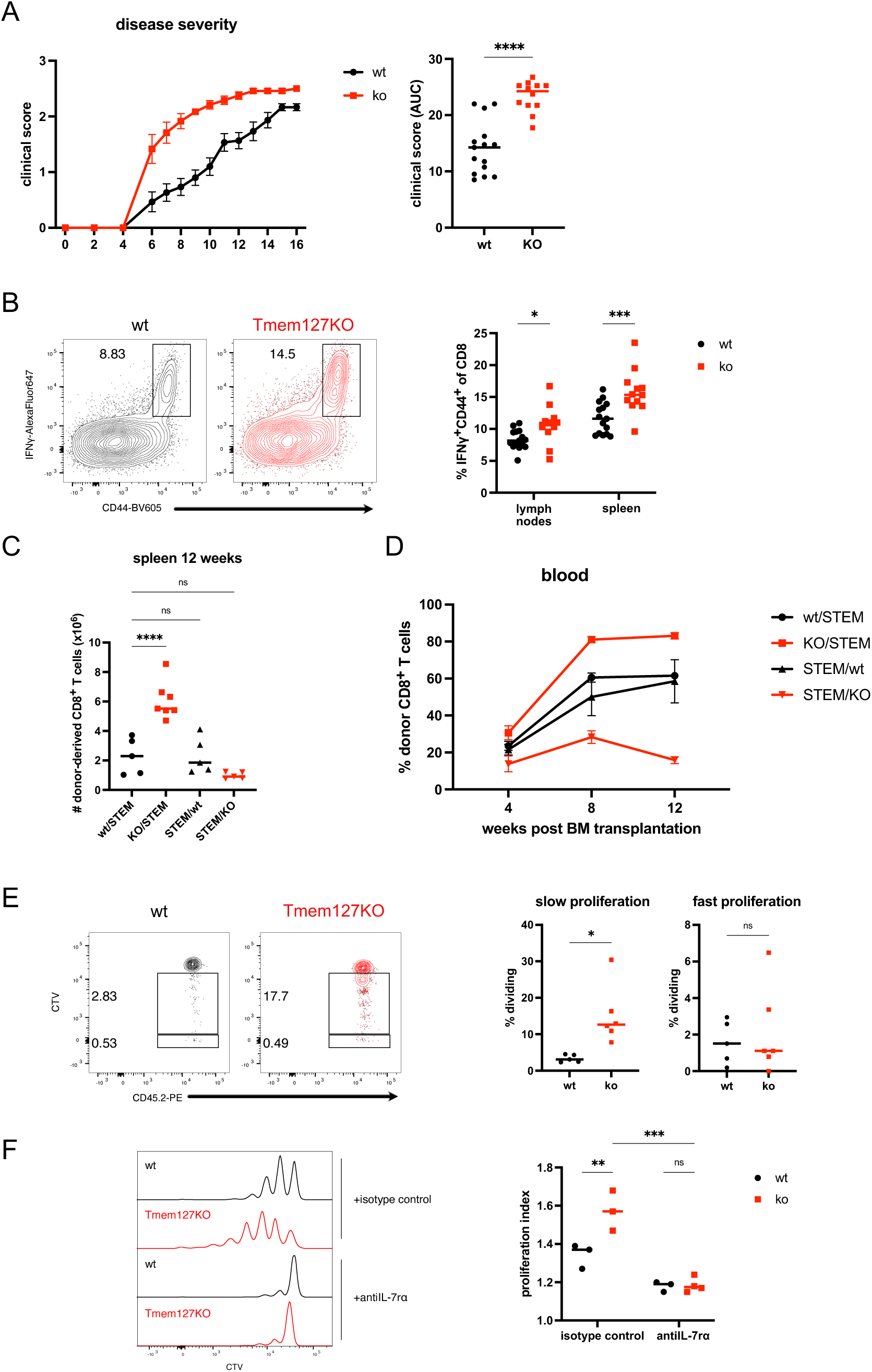
Tmem127 regulates homeostatic proliferation in vivo. **(A)** Clinical scores in MOG_35-55_-induced EAE model performed in Tmem127KO or wt mice, **(B)** Frequency of CD44^+^IFNγ^+^CD8^+^ T cells in lymphoid tissues in MOG_35-55_-induced EAE model preformed in Tmem127KO or wt mice, **(C)** Cell counts of donor-derived CD8^+^ T cells isolated from spleens of bone marrow chimeras 12 weeks after transplantation, **(D)** Frequency of donor-derived CD8^+^ T cells in blood of bone marrow chimeras, **(E)** Proliferation of donor naïve CD8^+^ T cells 14 days after adoptive transfer into immunocompetent STEM mice, **(F)** Proliferation of donor naïve CD8^+^ T cells 5 days after adoptive transfer into Rag1KO mice, injected together with anti-IL-7r blocking antibody or isotype control, dots represent mice and line median (or mean and SEM in A, left panel and D) of 2 independent experiments. P values were calculated using Mann-Whitney test (A), unpaired t-test (E), ordinary one-way ANOVA with multiple comparisons (C) or two-way ANOVA with Sidak’s multiple comparison test (B, F). ns, not significant; *p < 0.05; **p < 0.01; ***p < 0.001; ****p<0.0001.

To dissect the effects of Tmem127-deficiency on hematopoietic versus non-hematopoietic cells, we generated bone marrow chimeras since Tmem127KO mice harbor constitutive germline deletions. We transferred wt or Tmem127KO BM into sublethally irradiated CD45.1 congenic STEM mice (*39*) to investigate Tmem127’s hematopoietic cell-intrinsic role in reconstitution and development capacity. Vice versa, to investigate the role of Tmem127-deficiency in non-hematopoietic cells on immune reconstitution, we transferred CD45.1 STEM BM into wt or Tmem127KO recipients. 12 weeks after transplantation, control chimeras (wt -> STEM; STEM - > wt) displayed similar amounts of donor CD8^+^ T cells, while mice in the KO -> STEM group had an increased amount of donor-derived CD8^+^ T cells in the spleen (Fig. 5C). Transplantation of KO-> STEM also resulted in an increased frequency of donor CD8^+^ T cells compared to control groups (Fig. 5D). Vice versa, transplantation of control BM into Tmem127-deficient hosts resulted in a decreased frequency of donor CD8^+^ T cells (Fig. 5D). Thus, Tmem127-deficiency conferred a hematopoietic cell-intrinsic competitive advantage while Tmem127-deficiency in host cells constituted a competitive disadvantage to support wt cells. A similar trend was observed in donor BM hematopoietic stem cell frequencies with no difference in absolute cell numbers (fig. S5D). Finally, since many experiments pointed to an important role for T cell-intrinsic Tmem127 to regulate T cell homeostasis, we aimed to determine whether Tmem127KO T cells also cell-intrinsically hyperproliferated in vivo and if so, whether proliferation was IL-7-dependent. Of note, we hypothesized that γc hyperexpression could not only convey a competitive advantage to Tmem127KO T cells in lymphodepleted hosts but even in wt, lymphoreplete mice. Therefore, we adoptively transferred wt or Tmem127KO naïve CD8^+^ T cells (CD45.2^+^) (fig. S5E) to immunocompetent hosts (CD45.1^+^). Indeed, in analogy to the in vitro results, Tmem127KO T cells had an increased frequency of slow homeostatically proliferating cells (Fig. 5E). There was no difference, however, in fast proliferating cells (Fig. 5E), which start cycling in response to gut antigens and are activated by TCR (*40*). To test if the hyperproliferation was driven cell-intrinsically or extrinsically through endogenous host IL-7, we transferred T cells into Rag1KO mice. As in immunocompetent hosts, naive Tmem127KO T cells proliferated more than wt but cell proliferation from both genotypes was blocked by an IL-7 receptor-blocking antibody (Fig. 5F). This demonstrates that the driver of hyperpoliferation was extrinsic IL-7. In summary, Tmem127KO mice are more susceptible to EAE with an accelerated disease induction. Furthermore, γc hyperexpression caused by the absence of Tmem127 was functionally relevant in vivo and was the main driver of Tmem127KO T cell hyperproliferation.

## Discussion

Naïve T cells recirculate throughout the body in a quiescent state, ready to rapidly respond to cognate TCR/pMHC interaction to mount an immune response. They can remain in this dormant state for years (*41*), not unlike oocytes or stem cells. Naïve T cells, oocytes and stem cells employ posttranscriptional mechanisms to remain poised for rapid response (*41–43*). Upon activation, T cells rapidly alter their transcriptomes, miRNAs, proteomes and metabolism to enable cell growth, proliferation, differentiation and migration. Since many cellular decisions are guided by cell surface receptors, their expression needs to be tightly controlled. However, despite the central role of surface receptors in the interactions of T cells with their environment, our understanding of the steady-state regulation of membrane receptor expression levels is limited. It is known that even in the absence of agonistic TCR/pMHC interactions, naïve and memory T cells are not inactive. Instead, they dedicate significant resources to maintain a state of readiness, with a small fraction of cells entering the cell-cycle in the process of homeostatic proliferation (*2, 44, 45*). Starvation leads to impaired homeostatic proliferation of memory CD8 T cells even in conditions of severe T cell lymphopenia, supporting a role for active metabolism for T cell homeostasis (*46*). In addition, naïve T cell metabolism is controlled to maintain quiescence (*47, 48*). Btg1/2 proteins regulate the global mRNA stability in naïve T cells, including suppression of IL-7Rα and γc (*49*). Foxo1 is required for the maintenance of naïve T cells and, among various mechanisms, directly binds to the IL-7ra locus to induce IL-7Rα protein expression (*50, 51*). In contrast, Foxp1 is required to repress IL-7Rα by antagonizing Foxo1 and both transcription factors compete to regulate T cell quiescence (*52, 53*). Similar to our results, Foxp1-deficient and Btg1/2 double-deficient T cells proliferate in response to IL-7 without TCR stimulation in vitro (*49, 52*). Importantly, adoptively transferred Foxp1-deficient T cells upregulated IL-7Rα and proliferated even in lympho-replete recipient mice. Hyperproliferation also occurred in H-2K^b^- and H-2D^b^-deficient recipients but was blocked by IL-7 or IL-7R blocking antibodies, demonstrating that IL-7 and not TCR stimulation was the driving stimulus (*52*). In summary, the naïve T cell state is actively maintained (*44*) and among the known proteins that actively maintain T cell quiescence, several converge on the regulation of IL-7Rα, a known limiting factor for T cell homeostasis (*49–52*). Premature or inappropriate IL-7R expression can therefore alter T cell homeostasis. This is reminiscent of inappropriate Rorα expression by T cells lacking miR-17∼92 which results in impaired T_FH_ differentiation due to insufficient repression of alternative differentiation programs (*54*). Here, we shed light on a new mechanism how the second component of the IL-7R, the γc is restrained. We demonstrate that a lack of the membrane protein Tmem127 results in inappropriate γc and IL-7Rα expression, TCR-independent hypersensitivity to IL-7 and consequently dysregulated homeostasis. Of note, Foxo1-deficient T cells have severely impaired IL-7Rα expression while γc expression was unaffected (*50*). Thus, specific mechanisms tightly control the expression of both IL-7R components to preserve T cell homeostasis. Importantly, it was recently demonstrated that T cell preparedness is achieved by constant synthesis and rapid turnover of key RNA and proteins associated with activation, cell-cycle progression and transcription factors that maintain cell identity, quiescence and homeostasis. Thus, resting T cells selectively employ high protein turnover to adjust their proteome for transitioning cell states (*41*). We propose that Tmem127 regulates the turnover of key cell surface proteins including MHC and γc (*24*). Such regulation is likely relevant for other cell types for which MHC and γc are important. It will be important to investigate if and to what extent the Tmem127/IL-7R, or more generally Tmem127/γc axis, and Tmem127/MHC are active in other cell types such as innate lymphoid cells (ILCs), dendritic cells (DCs), macrophages, B cells, and various T cell subsets (*55*). To facilitate cellular investigations, we have generated a conditional mouse strain harboring a loxP flanked Tmem127 gene which we are open to share with potential collaborators.

With regards to the molecular mechanism, it was recently demonstrated that the adaptor protein Tmem127 forms a multi-protein complex with Susd6 and Wwp2 that directs MHC I towards lysosomal degradation (*24*). Here, we show that naive T cells utilize this complex to regulate their responsiveness to IL-7 and the rate of homeostatic proliferation by targeting γc. Since Tmem127 regulates at least two proteins with major functions in the immune system (MHC, γc) it will be important to further elucidate molecular details. An AF3 model predicts a direct interaction between Tmem127, Susd6, Wwp2 and γc. It will be important to determine whether the protein complex includes other proteins. For instance, Lrp10-deficient mice were recently reported to phenocopy many of the features observed in Tmem127-deficient mice including dysregulated IL-7R expression, hypersensitivity to IL-7, increased CD8^+^ T cells with an altered CD4/CD8 ratio and increased IL-7-driven, TCR-independent homeostatic T cell proliferation (*56*). Furthermore, Lrp10 encodes a putative endocytic receptor that posttranslationally suppresses IL-7R. The authors demonstrate an interaction between IL-7Rα with Lrp10. The interaction depends on the Lrp10 transmembrane and intracellular domain and at least partially on the IL-7Rα extracellular domain. Therefore, the authors conclude that the data does not support a direct protein-protein interaction but rather suggests that the interaction between IL-7Rα and Lrp10 may be mediated by a shared protein complex (*56*). Intriguingly, Lrp10 was identified as a MHC I negative regulator in the same CRISPR screening that identified the Susd6/Tmem127/Wwp2 complex to regulate MHC I (*24*). This raises the possibility that Lrp10 may be part of the Tmem127 protein complex. However, further studies are needed to clarify which proteins are involved in this protein complex centered around Tmem127 and whether the same proteins are involved in the regulation of IL-7Rα, MHC and γc. Molecular details should be studied extracellularly but investigations are also warranted to identify intracellular proteins which mediate protein degradation through this complex. Very little is known about which domains mediate protein-protein interactions, complex assembly and target degradation. Wwp2 contains 4 WW domains, which interact with specific PPxY or LPxY motifs. Tmem127’s C-terminal PPxY motif interacts with Wwp2 (*32*) but Wwp2-deficient mice neither display a CD4/CD8 shift nor develop splenomegaly, representing a distinct and milder phenotype than Tmem127KO mice (*57*). Similarly, in our hands, CRISPR-mediated Wwp2 removal resulted in a weaker phenotype than targeting Tmem127, echoing the finding that Wwp2 ablation augmented HLA expression less than removing Susd6 or Tmem127KO (*24*). Simultaneous Wwp2 interaction with the Tmem127 PPxY motif and the Susd6 LPxY motif could stabilize the complex and an interaction between Susd6 and Wwp2 was reported (*24*). Additionally, Lrp10 also contains a PPxY motif, which suggests it too could interact with the WW domain of Wwp2 and be a part of the complex. Together, these findings suggest that besides Wwp2, additional E3 ubiquitin ligases may ubiquitinate Tmem127 targets. For instance, the Nedd4 family of ubiquitin ligases includes eight proteins in mice (*58*), and two of them (Itch, Wwp1) have a similar domain structure as Wwp2 and could functionally compensate for Wwp2 loss. In fact, Itch shares a high structural homology and forms a complex with Wwp2 (*59*). Furthermore, Itch and Wwp2 synergize since mice double deficient for both develop spontaneous autoimmunity and a bias toward Th2 whereas the individual KOs have distinct phenotypes (*59*). Whether dysregulated γc expression contributes to the Th2 bias has not been investigated. Moreover, Itch associates with Foxo1 (required for IL-7Rα expression) and promotes its ubiquitination and degradation (*60*). Another candidate, STUB1, was identified in the screen for HLA negative regulators, and represents an E3 ubiquitin ligase known to affect MHC-I expression through downregulation of Ifngr1. Biochemical experiments confirmed an interaction with Tmem127 and Susd6 but it was not investigated further (*24*). Of note, Ifngr1 was identified in our Bio-ID experiment to be in proximity of Tmem127. In summary, there are many indications that the recently described Susd6/Tmem127/Wwp2 complex may involve more proteins extracellularly and that different E3 ligases may be involved in mediating MHC and/or γc degradation. Due to the complexity, additional studies are required to determine the molecular components, their assembly, motifs and how substrate specificity is achieved in different cell types.

Our study illustrates the challenges of identifying the targets regulated by the Susd6/Tmem127/Wwp2 complex. Several upregulated proteins in the CRISPR screening experiments seem to represent secondary effects of increased IL-7 signaling rather than direct targets. This is evident for proteins that were upregulated in an IL-7 concentration-dependent manner in vitro (CD25, CTLA-4, EOMES) but were not identified in the Bio-ID or proteomics experiments. Even the most consistently upregulated marker in the CRISPR experiments, ICOS, may be a secondary effect of IL-7 signaling since IL-7 is known to induce ICOS expression in NKT cells (*61*). Additionally, not all proteins found to be in proximity of Tmem127 are regulated by it, likely reflecting normal cell trafficking of the protein. However, our comparison of the Bio-ID results from a cell line and primary T cells shows that the Tmem127 targetome is likely unique in each cell type and depends on the protein expression profile of each cell type. Therefore, the physiological role and primary targets of Tmem127 regulation in other immune cell types might differ and our Tmem127 conditional mouse strain will facilitate further investigations.

Finally, our results clearly demonstrate that Tmem127 modulates disease severity. However, while Tmem127’s role in homeostatic proliferation is largely mediated by IL-7, its role in the pathogenesis of EAE is harder to ascribe to a single molecular mechanism. Dysregulated γc likely plays a role because IL-7 is known to aggravate autoimmunity and vice versa, γc or IL-7R blockade reduce the severity of EAE (*15, 62, 63*). However, Tmem127KO mice display a heavily altered proteome including increased MHC expression which could influence disease pathogenesis. Nevertheless, Tmem127 and its associated proteins offer opportunities for future therapeutic interventions. Previous studies suggested that therapeutic antibodies aiming to disrupt the Susd6/Tmem127/Wwp2 complex may improve immunotherapies by reverting cancer immune evasion in “cold” tumors through restoring MHCI expression (*24*). Our results suggest that this strategy could synergistically also promote T cell function and long-term maintenance by increasing γc and IL-7R. Furthermore, increased IL-7 signaling is well known to improve the efficiency of adoptive cell therapy (ACT) (*64*). The commonly used lymphodepletion prior to ACT reduces the cytokine sink created by the host cells, i.e. competition for IL-7 and IL-15 between endogenous and adoptively transferred cells (*64*). Therefore, deleting Tmem127, Susd6 or other proteins of this complex could enhance the function and competitiveness of adoptively transferred T cells, e.g. tumor infiltrating lymphocytes (TIL) or CAR T cells. This hypothesis is supported by our findings that Tmem127KO T cells and BM cells outcompeted their wildtype counterparts. Therefore, Tmem127KO T cells may have a competitive advantage or may reduce the need for lymphodepletion since they proliferated even in lymphoreplete hosts. Importantly, compared to wildtype controls, Lrp10 KO TILs displayed increased tumor infiltration, control of an autologous immunogenic tumor, increased IL-7R expression in the tumor microenvironment, resistance to T cell exhaustion and synergistic effects with anti-PD-1 immunotherapy (*56*). If Lrp10 is indeed part of the Susd6/Tmem127/Wwp2 complex, then the improved tumor control could be directly linked to MHC and/or γc expression. If not, the results nevertheless demonstrate the benefits of enhanced IL-7R expression on TILs for tumor control, as previously described (*38*). Vice versa, the role of Tmem127 in IL-7 signaling also suggests that the opposite strategy – enhancing Susd6/Tmem127/Wwp2 complex activity – could be used to treat autoimmunity or lymphoid malignancies that rely on IL-7 for their growth (*65, 66*). In summary, we uncovered Tmem127 as a new regulator of the γc. It’s remarkable that Tmem127 regulates two major immune proteins, MHC and γc, through a multi-protein complex with at least shared components.

## Acknowledgments

STEM mice were a kind gift of Jürg Schwaller. We thank Alessandro Dell’Aglio for assistance with animal experiments; the members of M.H.’s Doctoral Committee (Anne Spang and Petr Broz) and all members of the Jekerlab for critical feedback and suggestions throughout the project; Thomas Barthlott and Jessica Zuin for critical feedback on the manuscript. We acknowledge the following research core facilities at the University of Basel and the Department of Biomedicine: the animal facility for animal husbandry, the flow cytometry core facility for support, the Proteomics Core Facility (Katarzyna Buczak, Biozentrum, Basel) for proteomics; the bioinformatics core facility Basel for RNAseq data analysis; calculations were performed at sciCORE (http://scicore.unibas.ch/) scientific computing center at University of Basel; genomics facility Basel (ETH Zürich) for performing RNAseq. We acknowledge Natasa Savic from ETH Phenomics Center (ETH Zürich) for the generation of Tmem127KO mice; the technical support of the Core Facility Metabolomics and Proteomics (CF-MPC) at Helmholtz Munich and Juliane Merl-Pham for helping with the proteomics analysis of the Bio-ID experiments.

## Author contributions

Conceptualization: M.H., L.T.J.

Methodology: M.H., A.N., V.H., D.S., R.M, L.S., G.J., C.S.

Investigation: M.H., A.N., C.S., L.S., G.J., R.M., A.M.B.

Visualization: M.H., A.N., D.S., G.J., C.S.

Funding acquisition: L.T.J., V.H.

Project administration: M.H., L.T.J.

Supervision: L.T.J., V.H., A.K.P., S.H., M.B.

Writing – original draft: M.H., L.T.J.

Writing – review & editing: M.H., A.N., D.S., R.M., G.J., L.S., C.S., A.M.B., M.B., A.K.P., S.H., V.H., L.T.J.

## Data and materials availability

The Mass spectrometry data has been deposited in the ProteomeXchange Consortium via the MassIVE data repository (http://massive.ucsd.edu/) with the following accession numbers: PXD062457 (ProteomeXchange) and MSV000097467 (Massive). Bio-ID data was deposited to MassIVE repository under accession number MSV000097926. RNA sequencing data can be viewed at Gene Expression Omnibus (GEO) under accession number GSE293773. The TCR sequencing data have been deposited at the European Nucleotide Archive (ENA) under the accession number PRJEB89688. Tmem127KO mice (strain 1) are available upon reasonable request. Tmem127KO mouse strains 2-5 are available as cryopreserved sperm upon reasonable request.

## Supplementary Materials

Materials and Methods

Figs. S1 to S5

Tables S1 to S2

References (*67–82*)

## References

1. I. den Braber et al., Maintenance of peripheral naive T cells is sustained by thymus output in mice but not humans. Immunity 36, 288–297 (2012).

2. I. Bains, R. Antia, R. Callard, A. J. Yates, Quantifying the development of the peripheral naive CD4+ T-cell pool in humans. Blood 113, 5480–5487 (2009).

3. K. S. Schluns, W. C. Kieper, S. C. Jameson, L. Lefrancois, Interleukin-7 mediates the homeostasis of naive and memory CD8 T cells in vivo. Nat Immunol 1, 426–432 (2000).

4. J. T. Tan et al., IL-7 is critical for homeostatic proliferation and survival of naive T cells. Proc Natl Acad Sci U S A 98, 8732–8737 (2001).

5. A. W. Goldrath, M. J. Bevan, Low-affinity ligands for the TCR drive proliferation of mature CD8+ T cells in lymphopenic hosts. Immunity 11, 183–190 (1999).

6. J. H. Park et al., Suppression of IL7Ralpha transcription by IL-7 and other prosurvival cytokines: a novel mechanism for maximizing IL-7-dependent T cell survival. Immunity 21, 289–302 (2004).

7. L. Swainson, E. Verhoeyen, F. L. Cosset, N. Taylor, IL-7R alpha gene expression is inversely correlated with cell cycle progression in IL-7-stimulated T lymphocytes. J Immunol 176, 6702–6708 (2006).

8. C. M. Henriques, J. Rino, R. J. Nibbs, G. J. Graham, J. T. Barata, IL-7 induces rapid clathrin-mediated internalization and JAK3-dependent degradation of IL-7Ralpha in T cells. Blood 115, 3269–3277 (2010).

9. N. Sauvonnet, A. Dujeancourt, A. Dautry-Varsat, Cortactin and dynamin are required for the clathrin-independent endocytosis of gammac cytokine receptor. J Cell Biol 168, 155–163 (2005).

10. S. M. Russell et al., Interaction of IL-2R beta and gamma c chains with Jak1 and Jak3: implications for XSCID and XCID. Science 266, 1042–1045 (1994).

11. T. Miyazaki et al., Functional activation of Jak1 and Jak3 by selective association with IL-2 receptor subunits. Science 266, 1045–1047 (1994).

12. B. M. Foxwell, C. Beadling, D. Guschin, I. Kerr, D. Cantrell, Interleukin-7 can induce the activation of Jak 1, Jak 3 and STAT 5 proteins in murine T cells. Eur J Immunol 25, 3041–3046 (1995).

13. A. V. Villarino, et al., A central role for STAT5 in the transcriptional programing of T helper cell metabolism. Sci Immunol 7, eabl9467 (2022).

14. M. Noguchi et al., Interleukin-2 receptor gamma chain mutation results in X-linked severe combined immunodeficiency in humans. Cell 73, 147–157 (1993).

15. A. Le Floc’h et al., Blocking common gamma chain cytokine signaling ameliorates T cell-mediated pathogenesis in disease models. Sci Transl Med 15, eabo0205 (2023).

16. D. Baumjohann, Diverse functions of miR-17-92 cluster microRNAs in T helper cells. Cancer Lett 423, 147–152 (2018).

17. C. Xiao et al., Lymphoproliferative disease and autoimmunity in mice with increased miR-17-92 expression in lymphocytes. Nat Immunol 9, 405–414 (2008).

18. L. He et al., A microRNA polycistron as a potential human oncogene. Nature 435, 828–833 (2005).

19. V. Olive et al., miR-19 is a key oncogenic component of mir-17-92. Genes Dev 23, 2839–2849 (2009).

20. Y. C. Han et al., An allelic series of miR-17 approximately 92-mutant mice uncovers functional specialization and cooperation among members of a microRNA polycistron. Nat Genet 47, 766–775 (2015).

21. M. Regelin et al., Responsiveness of Developing T Cells to IL-7 Signals Is Sustained by miR-17 approximately 92. J Immunol 195, 4832–4840 (2015).

22. K. Liao et al., Critical roles of the miR-17 approximately 92 family in thymocyte development, leukemogenesis, and autoimmunity. Cell Rep 43, 114261 (2024).

23. M. Dolz et al., Forced expression of the non-coding RNA miR-17 approximately 92 restores activation and function in CD28-deficient CD4(+) T cells. iScience 25, 105372 (2022).

24. X. Chen et al., A membrane-associated MHC-I inhibitory axis for cancer immune evasion. Cell 186, 3903–3920 e3921 (2023).

25. E. K. Brinkman, T. Chen, M. Amendola, B. van Steensel, Easy quantitative assessment of genome editing by sequence trace decomposition. Nucleic Acids Res 42, e168 (2014).

26. J. G. Doench et al., Optimized sgRNA design to maximize activity and minimize off-target effects of CRISPR-Cas9. Nat Biotechnol 34, 184–191 (2016).

27. G. Katz et al., T cell receptor stimulation impairs IL-7 receptor signaling by inducing expression of the microRNA miR-17 to target Janus kinase 1. Sci Signal 7, ra83 (2014).

28. J. Irie-Sasaki et al., CD45 is a JAK phosphatase and negatively regulates cytokine receptor signalling. Nature 409, 349–354 (2001).

29. J. H. Cho et al., CD45-mediated control of TCR tuning in naive and memory CD8(+) T cells. Nat Commun 7, 13373 (2016).

30. A. H. Courtney et al., CD45 functions as a signaling gatekeeper in T cells. Sci Signal 12, (2019).

31. S. Srikantan et al., The tumor suppressor TMEM127 regulates insulin sensitivity in a tissue-specific manner. Nat Commun 10, 4720 (2019).

32. E. Alix et al., The Tumour Suppressor TMEM127 Is a Nedd4-Family E3 Ligase Adaptor Required by Salmonella SteD to Ubiquitinate and Degrade MHC Class II Molecules. Cell Host Microbe 28, 54–68 e57 (2020).

33. K. P. Hoefig et al., Defining the RBPome of primary T helper cells to elucidate higher-order Roquin-mediated mRNA regulation. Nat Commun 12, 5208 (2021).

34. J. Abramson et al., Accurate structure prediction of biomolecular interactions with AlphaFold 3. Nature 630, 493–500 (2024).

35. A. J. M. Howden et al., Quantitative analysis of T cell proteomes and environmental sensors during T cell differentiation. Nat Immunol 20, 1542–1554 (2019).

36. M. Harndahl, M. Rasmussen, G. Roder, S. Buus, Real-time, high-throughput measurements of peptide-MHC-I dissociation using a scintillation proximity assay. J Immunol Methods 374, 5–12 (2011).

37. X. T. Cheng et al., Characterization of LAMP1-labeled nondegradative lysosomal and endocytic compartments in neurons. J Cell Biol 217, 3127–3139 (2018).

38. C. L. Mackall, T. J. Fry, R. E. Gress, Harnessing the biology of IL-7 for therapeutic application. Nat Rev Immunol 11, 330–342 (2011).

39. F. E. Mercier, D. B. Sykes, D. T. Scadden, Single Targeted Exon Mutation Creates a True Congenic Mouse for Competitive Hematopoietic Stem Cell Transplantation: The C57BL/6-CD45.1(STEM) Mouse. Stem Cell Reports 6, 985–992 (2016).

40. W. C. Kieper et al., Recent immune status determines the source of antigens that drive homeostatic T cell expansion. J Immunol 174, 3158–3163 (2005).

41. T. Wolf et al., Dynamics in protein translation sustaining T cell preparedness. Nat Immunol 21, 927–937 (2020).

42. J. D. Vassalli, A. Stutz, Translational control. Awakening dormant mRNAs. Curr Biol 5, 476–479 (1995).

43. T. H. Cheung, T. A. Rando, Molecular regulation of stem cell quiescence. Nat Rev Mol Cell Biol 14, 329–340 (2013).

44. J. Sprent, C. D. Surh, Normal T cell homeostasis: the conversion of naive cells into memory-phenotype cells. Nat Immunol 12, 478–484 (2011).

45. D. Tzachanis et al., Tob is a negative regulator of activation that is expressed in anergic and quiescent T cells. Nat Immunol 2, 1174–1182 (2001).

46. S. S. Iyer et al., Protein energy malnutrition impairs homeostatic proliferation of memory CD8 T cells. J Immunol 188, 77–84 (2012).

47. K. Yang, G. Neale, D. R. Green, W. He, H. Chi, The tumor suppressor Tsc1 enforces quiescence of naive T cells to promote immune homeostasis and function. Nat Immunol 12, 888–897 (2011).

48. N. M. Chapman, M. R. Boothby, H. Chi, Metabolic coordination of T cell quiescence and activation. Nat Rev Immunol 20, 55–70 (2020).

49. S. S. Hwang et al., mRNA destabilization by BTG1 and BTG2 maintains T cell quiescence. Science 367, 1255–1260 (2020).

50. Y. M. Kerdiles et al., Foxo1 links homing and survival of naive T cells by regulating L-selectin, CCR7 and interleukin 7 receptor. Nat Immunol 10, 176–184 (2009).

51. W. Ouyang, O. Beckett, R. A. Flavell, M. O. Li, An essential role of the Forkhead-box transcription factor Foxo1 in control of T cell homeostasis and tolerance. Immunity 30, 358–371 (2009).

52. X. Feng et al., Transcription factor Foxp1 exerts essential cell-intrinsic regulation of the quiescence of naive T cells. Nat Immunol 12, 544–550 (2011).

53. C. N. Skon, S. C. Jameson, Fox factors fight over T cell quiescence. Nat Immunol 12, 522–524 (2011).

54. D. Baumjohann et al., The microRNA cluster miR-17 approximately 92 promotes TFH cell differentiation and represses subset-inappropriate gene expression. Nat Immunol 14, 840–848 (2013).

55. Y. Rochman, R. Spolski, W. J. Leonard, New insights into the regulation of T cells by gamma(c) family cytokines. Nat Rev Immunol 9, 480–490 (2009).

56. J. Russell et al., Lrp10 suppresses IL7R limiting CD8 T cell homeostatic expansion and anti-tumor immunity. EMBO Rep 25, 3601–3626 (2024).

57. Y. Yang et al., E3 ligase WWP2 negatively regulates TLR3-mediated innate immune response by targeting TRIF for ubiquitination and degradation. Proc Natl Acad Sci U S A 110, 5115–5120 (2013).

58. R. J. Ingham, G. Gish, T. Pawson, The Nedd4 family of E3 ubiquitin ligases: functional diversity within a common modular architecture. Oncogene 23, 1972–1984 (2004).

59. D. Aki et al., The E3 ligases Itch and WWP2 cooperate to limit T(H)2 differentiation by enhancing signaling through the TCR. Nat Immunol 19, 766–775 (2018).

60. N. Xiao et al., The E3 ubiquitin ligase Itch is required for the differentiation of follicular helper T cells. Nat Immunol 15, 657–666 (2014).

61. A. J. Lee et al., Regulation of natural killer T-cell development by deubiquitinase CYLD. EMBO J 29, 1600–1612 (2010).

62. H. Winer et al., IL-7: Comprehensive review. Cytokine 160, 156049 (2022).

63. J. J. Ashbaugh et al., IL7Ralpha contributes to experimental autoimmune encephalomyelitis through altered T cell responses and nonhematopoietic cell lineages. J Immunol 190, 4525–4534 (2013).

64. L. Gattinoni et al., Removal of homeostatic cytokine sinks by lymphodepletion enhances the efficacy of adoptively transferred tumor-specific CD8+ T cells. J Exp Med 202, 907–912 (2005).

65. I. Lodewijckx, J. Cools, Deregulation of the Interleukin-7 Signaling Pathway in Lymphoid Malignancies. Pharmaceuticals (Basel) 14, (2021).

66. H. Dooms, Interleukin-7: Fuel for the autoimmune attack. J Autoimmun 45, 40–48 (2013).

