## Supplementary material for "Tmem127-mediated immune receptor degradation regulates T cell homeostasis through the common gamma chain": Suppl. Material

**The PDF file includes:**

Materials and Methods
Figs. S1 to S5
References

**Other Supplementary Materials for this manuscript include the following:**

Tables S1 to S2

#### Materials and Methods

Screening design and data analysis

Three guide RNAs targeting each of 68 genes were distributed randomly between 3 96-well plates containing control samples: PtprcKO, TracKO, PtenKO, and non-targeting controls (NTC): CD28KO and wt. Control samples were placed in the same positions between replicate plates. To reduce the plate effect, all gRNAs targeting the same gene were always allocated to the same plate. After data acquisition, plate effects were further controlled by the scaling normalization method. Except for NTC_CD28KO, which was present in triplicate on each plate, others were present in duplicates. The first step was performed per plate, and then the plates were merged at the end. Raw MFI and percentage features of the panels were log-transformed using R package proBatch (*67*). To correct the batch effect using the control information, we decided to set the same value ranges according to controls and then shift panel distribution values. Within each plate, we computed the mean of control values (duplicates/triplicates) for each of the panel features independently. Then, we selected minimum and maximum values and, to combine plates, a consensus approach was used: when two of the three plates agreed, min and max values for a given feature were computed between the agreed plates. When all plates disagreed, we arbitrarily set the minimum value of NTC_CD28KO control and the maximum value of NTC_WT.

Normalized values distributions were then shifted within min and max values of controls defined by a consensus approach, within each plate. Plates were finally merged and scaled (z-score). Principal component analysis (PCA) was performed on batch-corrected data. We performed a (PCA) limited to the first 10 principal components (PCs) using the pca function (method = “nipals” and center = T) from R package pcaMethods (*68*).

Mice

C57BL/6N mice were obtained from the Charles River Laboratory (stock number 027). Tmem127KO lines were created by the ETH Zurich EPIC facility, using gRNA1. B6.129S2-Cd28tm1Mak/J (CD28KO) and B6.129S7-Rag1tm1Mom/J (Rag1KO) mice were obtained from SwimMR. C57BL/6N-CD45.1$^{STEM}$ (STEM) mice were a gift from Jürg Schwaller. $Gt(ROSA)26Sor^{tm1(rtTA*M2)Jae}$ mice (C57BL/6 background) were purchased from The Jackson Laboratory and housed at the Ludwig-Maximilians-University Munich (LMU) in a specific-pathogen-free (SPF) barrier facility under a 12 h/12 h dark/light cycle at 20–24°C and 45–65% humidity in accordance with the LMU institutional, state and federal guidelines. These mice were used for doxycycline-inducible expression experiments.

All other mice were kept in specific pathogen-free conditions in ventilated cages in rooms with artificial light with daylight spectra, 12 hours light/12 hours dark with dusk and dawn phases of 30 minutes, maximum 200 lux. The temperature in the rooms was between 20°C and 24°C, and humidity was 45-65%. All mice were fed a standard facility diet.

Most in vitro experiments and immunophenotyping of Tmem127KO mice were performed on groups containing a 1:1 ratio of male and female replicates. RNA sequencing and proteomics were performed on female mice. Adoptive transfer experiments were performed on male mice. Bone marrow chimeras and EAE experiments were performed with female mice. All experiments were performed on cells isolated from 6-20 weeks old mice, age-matched in each experiment.

All animal experiments were done in compliance with Swiss federal and cantonal laws and were approved by the Animal Research Commission of Canton of Basel-Stadt, Switzerland.

Naïve T cell isolation, in vitro activation, and homeostatic proliferation assays

Naïve $CD4^+$ and $CD8^+$ T cells were isolated using corresponding STEMCELL kits (19765A, 19858A) according to the manufacturer's instructions.

For in vitro homeostatic proliferation assays, cells were cultured for 4 (naïve $CD8^+$) or 6 (CRISPR-edited naïve $CD4^+$ and naïve $CD8^+$) days in vitro in T cell medium supplemented with recombinant mouse IL-7 (R&D, 407-ML-005) at concentrations 1 or 5 ng/ml.

For in vitro activation, flat-bottom 96-well plates (Corning) were coated overnight at 4°C with PBS dilution of anti-CD3/anti-CD28 (0.5/0.2 or 5/2 μg/ml) antibodies (Bio X Cell, BP0001-1, BE0015-5), and washed with PBS once, cells were cultured on coated plates for 2 or 3 days.

Cells were cultured in the incubator at a temperature of 37°C and 5% $CO_2$ content.

Cas9 ribonucleoprotein electroporation in naïve mouse T cells

gRNAs were prepared by annealing crRNAs (IDT) with tracrRNA (IDT) in a 1:1 molar ratio at 95°C for 5 minutes. 0.8x volume of 100 mg/ml poly-L-glutamic acid (PGA, Sigma) solution was mixed with 1 volume of gRNA solution. CRISPR Cas9 ribonucleoproteins were prepared by mixing gRNA/PGA solution and recombinant Cas9 (Berkley) in a 3:1 molar ratio. $6 \times 10^5$- $1 \times 10^7$ naïve $CD4^+$ or $CD8^+$ T cells were washed with PBS and resuspended in P4 electroporation buffer (Lonza) in a final volume of 20 μl per electroporation, mixed with RNP solution and electroporated in Nucleofector (Lonza) with pulse DS137. Knockout efficiency was assessed by comparing Sanger sequencing (Microsynth) data from edited and non-targeting control samples. DNA for Sanger sequencing was isolated using 30 μl of QuickExtract solution (Lucigen) per sample ($6*10^5$ cells).

Flow cytometry staining

For viability staining, cells were stained with eFluor780 (Invitrogen, 65-0865-14) according to the manufacturer's instructions. For surface staining, $1 \times 10^5$- $3 \times 10^6$ cells were incubated in 50 μl antibody solution in FACS buffer for 30 min at 4 °C, centrifuged (370 g, 4 min, 4°C) with 150 μl FACS buffer, and fixed with fixation buffer (Invitrogen, 008222-49) for 30 min at 4°C. For intracellular staining, cells were permeabilized with 100 μl fixation/permeabilization buffer (Invitrogen, 005123-42, 005123-56) for 15 min at RT, centrifuged (370 g, 4 min, 4°C) with permeabilization buffer (Invitrogen, 00-8333-56) and incubated in 50 μl antibody solution in permeabilization buffer for 1 hour at RT. For phospho-STAT5 staining, $3 \times 10^6$ naïve $CD8^+$ T cells were cultured overnight with 5 ng/ml IL-7, fixed with 4% paraformaldehyde (Sigma-Aldrich, 16005-1KG-R) solution for 10 min at RT, and permeabilized with methanol (Sigma-Aldrich, 34860-1L-R) on ice for 30 min.

Retroviral transduction of MEF cells and primary mouse T cells

HEK293T cells, pre-treated with 25 μM chloroquine, were co-transfected with 5 μg of the packaging vector pCL-Eco (Addgene #12371) and 50 μg of the respective expression plasmids using calcium phosphate transfection. After 4–6 hours, the medium was replaced with fresh medium, and the cells were cultured for an additional 40 hours. The viral particle-containing supernatant was collected, filtered through a 0.45 μm filter, and supplemented with polybrene (10 μg/mL) for subsequent transduction.

Retroviral transduction of rtTA-positive $CD8^+$ T cells was performed during T cell activation. Primary murine $CD8^+$ T cells were isolated from single-cell suspensions prepared from spleens and peripheral lymph nodes. Cells were resuspended in T cell isolation buffer (PBS supplemented with 2% FCS and 1 mM EDTA) and erythrocytes were lysed using TAC lysis buffer (13 mM Tris, 140 mM NH4Cl, pH 7.2). $CD8^+$ T cells were then isolated by negative selection using the EasySep™ Mouse $CD8^+$ T Cell Isolation Kit (STEMCELL). A total of 2.5–3 × $10^6$ T cells were activated in vitro in 6-well plates with plate-bound anti-CD3 (0.1 µg/mL; clone 2C11, in-house) and anti-CD28 antibodies (1.0 µg/mL; clone 37.5 N, in-house), cross-linked with rabbit anti-hamster IgG (anprotec, #AC-AR-0034). Cells were cultured in T cell medium (DMEM (Gibco) supplemented with 100 U/mL Pen-Strep (Gibco), 10 mM HEPES (Gibco), 50 µM β-mercaptoethanol (Gibco), and 1x nonessential amino acids (Gibco) and 10% FCS (Bio&SELL). After 40 hours of activation, the supernatant from the activated plate-bound $CD8^+$ T cells was replaced with the virus-containing supernatant. Spin-inoculation was conducted at 860g for 1 hour at 18°C. The $CD8^+$ T cells were then incubated at 37°C and 5% $CO_2$ for 4–6 hours before being cultured in fresh medium supplemented with 200 U/mL human IL-2 (Novartis). On the third day post-transduction, doxycycline was added to induce expression from the pRetroXTight plasmid, and the cells were harvested 22–24 hours later.

Retroviral transduction of rtTA-positive MEF cells was carried out the day after seeding. The culture supernatant was replaced with virus-containing supernatant, followed by spin-inoculation at 300g for 2 hours at 32°C. Cells were then incubated for 4–6 hours at 37°C and 10% $CO_2$. The viral medium was replaced with a fresh medium, and the cells were expanded on day 2 post-transduction. Cells were seeded again on day 4 post-transduction and treated with doxycycline the following day. MEF cells for confocal microscopy were analyzed 18 hours later.

RNA-seq

Sequencing was performed at Genomics Facility Basel (ETH Zurich). All naïve $CD8^+$ T cells isolated from spleens of wt and Tmem127KO animals were lysed and RNA isolated using a Quick-RNA MiniPrep kit (Zymo research, R1055). Quality control was performed using TapeStation (Agilent). The library was prepared using the Illumina TruSeq stranded mRNA library preparation kit. Sequencing was performed on an Illumina NovaSeq 6000 instrument and paired-end 51 bp reads were produced. Read quality was assessed with the FastQC tool (version 0.11.5). Reads were mapped to the mouse genome mm39 with STAR (version 2.7.10a) (*69*) with default parameters, except filtering out multimapping reads with more than 10 alignment locations (outFilterMultimapNmax=10) and filtering reads without evidence in the spliced junction table (outFilterType="BySJout"). The featureCounts (*70*) function from the Rsubread package (version 2.0.6) was used to count the number of reads (5' ends) overlapping with the exons of each gene (Ensembl release 110), assuming an exon union model. All subsequent analyses were performed using the R software (version 4.3.1). The Bioconductor package edgeR (version 3.42.4) (*71*) was used for differential gene expression analysis. Between samples, normalization was done using the TMM method (*72*). Only genes with CPM (counts per million reads mapped) values more than 1 in at least 3 samples were retained. Principal component analysis (PCA) was performed on the log-transformed normalized CPM values. The quasi-likelihood testing framework (function glmQLFit with option legacy=FALSE and function glmQLFTest) was used to perform comparisons between genotypes. P-values were adjusted by controlling the false discovery rate (FDR; Benjamini-Hochberg method) and genes with an FDR lower than 5% were considered significant. Transcription factors activity was inferred using the decoupleR package (version 2.6.0) (*73*). As input, we used the CollecTRI collection, including a curated collection of TFs and their targets compiled from multiple resources, accessed using the OmnipathR package (version 3.8.2) (*74*).

TCR-seq

DNA was isolated from naïve CD4$^+$ and naïve CD8$^+$ T cells of wt and Tmem127KO animals using quickDNA miniprep plus kit (Zymo research, D4069). Amplification of the rearranged V, D, and J gene segments of the murine TRB locus was performed in a multiplex PCR with 250-500 ng genomic DNA template and a murine TRB primer set (*75*). After barcoding and quality control, the TRB libraries were sequenced and demultiplexed on an Illumina MiSeq sequencer (600-cycle single-index, paired-end run, V3 chemistry). Read alignment was performed using the MiXCR framework (*76*) and the default reference library. Nonproductive reads and sequences with less than two reads were not included in downstream bioinformatics analyses. All repertoires were proportionally normalized to 30,000 reads. Each unique complementarity-determining region 3 (CDR3) nucleotide sequence was a clone. All analyses were performed using RStudio (version 1.1.456) and the tcR package as described (*75*).

Proteomics

After washing in PBS, a pellet containing $10^6$ naïve CD8$^+$ T cells was frozen in liquid nitrogen, or cells were activated in vitro for 72 hours and frozen afterwards. Cell pellets were thawed and lysed in 50 µL of lysis buffer (1% Sodium deoxycholate (SDC), 10 mM TCEP, 100 mM Tris, pH=8.5) using twenty cycles of sonication (30 s on, 30 s off per cycle) on a Bioruptor (Dianode). Following sonication, proteins in the cell lysates were reduced by TCEP at 95°C for 10 min. Proteins were alkylated using 15 mM chloroacetamide at 37°C for 30 min and further digested using sequencing-grade modified trypsin (1/50 w/w, ratio trypsin/protein; Promega, USA) at 37°C for 12 hours. After digestion, the samples were acidified using TFA (final 1%). Peptide desalting was performed using iST cartridges (PreOmics, Germany) following the manufacturer's instructions. After drying the samples under vacuum and storing at -20°C, enriched peptides were resuspended in 0.1% aqueous formic acid and subjected to LC-MS/MS analysis using an Orbitrap Eclipse Mass Spectrometer fitted with a Vanquid Neo nanoLC (both Thermo Fisher Scientific) and a custom-made column heater set to 60°C. Peptides were resolved using an RP-HPLC column (75µm × 30cm) packed in-house with C18 resin (ReproSil-Pur C18–AQ, 1.9 µm resin; Dr. Maisch GmbH) at a flow rate of 0.2 µLmin-1. The following gradient was used for peptide separation: from 4% B to 10% B over 5 min, to 35% B over 45 min to 50% B over 10 min to 95% B over 1 min followed by 10 min at 95% B. Buffer A was 0.1% formic acid in water and buffer B was 80% acetonitrile, 0.1% formic acid in water.

The mass spectrometer was operated in DIA mode with a cycle time of 3 seconds. MS1 scans were acquired in the Orbitrap in centroid mode at a resolution of 120,000 FWHM (at 200 m/z), a scan range from 390 to 910 m/z, normalized AGC target set to 300%, and maximum ion injection time mode set to Auto. MS2 scans were acquired in the Orbitrap in centroid mode at a resolution of 15,000 FWHM (at 200 m/z), precursor mass range of 400 to 900, quadrupole isolation window of 10 m/z with 1 m/z window overlap, a defined first mass of 120 m/z, normalized AGC target set to 800% and a maximum injection time of 22 ms. Peptides were fragmented by HCD with collision energy set to 28% and one microscan was acquired for each spectrum.

The acquired files were searched using the Spectronaut (Biognosys v17.6) directDIA workflow using standard settings. The search was done against a Mus Musculus database (downloaded from Uniprot on 20220222). Quantitative fragment ion data (F.Area) was exported from Spectronaut and analyzed using the MSstats R package (version 4.8.2) (*77*). Data was normalized using the default normalization option "equalizedMedians", imputed using "AFT model-based imputation" and p-values and q-values for pairwise comparisons were calculated using the default settings of the MSstats package.

Cell line culture

HEK293T and MEFs were maintained in DMEM (Gibco) supplemented with 10% (v/v) FCS (Bio&SELL), 100 U/mL Pen-Strep (Gibco) and 10 mM HEPES (Gibco). Cells were cultured at 37°C in an atmosphere containing 10% CO2. The cell lines were obtained from ATCC.

Expression plasmids

The full coding sequence (CDS) of murine Tmem127 isoform 1, along with its truncated versions, Il2rg variant A, or Wwp2, were cloned into the MSCV–IRES–Thy-1.1 expression vector. Additionally, the full CDS of Tmem127 and its truncated variants were inserted into the pRetro-Xtight expression plasmid (Clontech) under the control of a Tet-responsive promoter and fused at the C-terminus with a "GGSGGSGG" linker to either eGFP or a mutated version of biotin ligase (BirA*) from *E. coli*. The insertion of CDS sequences into the expression plasmids was facilitated using the Gateway® cloning system (Thermo Fisher Scientific). Fusion of the CDS with the linker-eGFP or linker-BirA* was carried out using a PCR product containing the sequence of interest with overhangs at the 5'-end to the Tmem127 CDS and at the 3'-end to the plasmid backbone, serving as a megaprimer in a PCR reaction following the method described in (*78*). The linker sequence was introduced in a subsequent step using the same approach. Primer sequences are available upon request.

Calcium phosphate transfection

HEK293T cells, pre-treated with 25 µM chloroquine, were co-transfected with 50 µg of each respective expression plasmid using calcium phosphate as the transfection reagent. After 4-6 hours, the medium was replaced with a fresh culture medium. The following day, the cells were harvested, washed with ice-cold PBS, and the cell pellet was snap-frozen in liquid nitrogen before being stored at -80°C.

Bio-ID

The proximity-dependent biotin identification assay was performed on rtTA-positive MEF cells or $CD8^+$ T cells.

For rtTA-positive MEF cells, each sample had 3 technical replicates of Tmem127-BirA*, Tmem127 C-terminal truncated-BirA*, or eGFP-BirA* transduced MEF cells with 50% confluency in 2x15cm dishes were prepared. Six hours after the addition of doxycycline, 50µM biotin was added for 18 h. Cells were trypsinized and washed twice with PBS. For rtTA $CD8^+$ T cells, each sample was performed in 3 biological replicates that were transduced with Tmem127-BirA*, Tmem127 C-terminal truncated-BirA*, or eGFP-BirA*. On day 3 after transduction at a density of 1mio/mL doxycycline was added, followed by 50µM biotin after 6h for another 16 h. Cells were washed twice with PBS.

The samples were processed as described in (*33*) with the minor change, that proteins were eluted from streptavidin beads in 30µL. For identification and quantification of proteins, samples were proteolysed by a modified filter-aided sample preparation as described (*79, 80*), and eluted peptides were analyzed by LC–MSMS on a QExactive HFX mass spectrometer (ThermoFisher Scientific) coupled directly to an UItimate 3000 RSLC nano-HPLC (Dionex) as described (*81*). Generated raw files were quantitatively analyzed in the Proteome discoverer 2.5 software (Thermo Scientific) for peptide and protein identification and quantification. Database search was performed using the Sequest HT search engine against the SwissProt Mouse database (Release

2020_02, 17061 sequences). Match-between runs for label-free quantification were limited to a retention time window of 1 minute and a mass shift of 0.5 ppm. Peptide abundance values were normalized on the total peptide amount. Missing values were imputed by low abundance resampling separately for each sample. Protein abundances were calculated as the average of the 3 most abundant unique peptide group intensity values (TOP3) with an XCorr score >1. Protein identifications and quantifications were exported and filtered for a protein false discovery rate <5%. Protein abundance ratios and Student's T-tests with Benjamini-Hochberg correction were calculated using the normalized and imputed protein abundance values.

Co-immunoprecipitation

Protein-A or Protein-G Dynabeads (40 µL; Invitrogen) were coupled to 2 µg of either anti-IL2rg antibody (CST, Cat# 18526) or GFP antibody (Merck, Cat# 11814460001) in lysis buffer (20 mM Tris-HCl, pH 7.5, 150 mM NaCl, 0.25% NP-40 and 1.5 mM $MgCl_2$) under constant rotation at 4°C overnight. T cells or HEK293T cells were washed with ice-cold PBS, snap-frozen in liquid nitrogen and stored at -80°C. Pellets were lysed in lysis buffer supplemented with 1 mM DTT, and 1x complete EDTA-free protease inhibitor cocktail (Roche) for 30 minutes on ice at 4°C. Lysates were clarified by centrifugation at 20,000×g for 10 minutes, and protein concentration was determined using a Bradford assay (Bio-Rad). The input fraction was denatured by heating at 95°C for 5 minutes in 4× Laemmli buffer (314 mM Tris, 50% glycerol, 5% SDS, 5% β-mercaptoethanol, and 0.01% bromophenol blue, pH 6.8).

The antibody-coupled beads were washed three times with lysis buffer and incubated with 1.5 mg of HEK293T protein lysate or 3.2–3.5 mg of $CD8^+$ T cell protein lysate in 500 µL of lysis buffer supplemented with 1 mM DTT, and 1x complete EDTA-free protease inhibitor cocktail (Roche) for 4 hours at 4°C under constant rotation. Beads were then washed three times with lysis buffer, resuspended in 25 µL of 1× Laemmli buffer, and boiled at 95°C for 10 minutes. A total of 25 µg of the input fraction and the entire 25 µL of the co-immunoprecipitated samples were loaded onto an SDS-PAGE gel, transferred to a methanol-activated PVDF membrane, and probed with the appropriate primary (anti-Il2rg, mouse anti-GFP or rabbit anti-GFP (Invitrogen, A-11122)) and HRP-conjugated secondary antibodies (CST). Signal detection was carried out using ECL western blotting reagents (either in-house produced ECL or SuperSignal™ West Femto, Thermo Fisher), and images were captured using the iBright 1500 system (Invitrogen).

Adoptive transfer of naïve $CD8^+$ cells

2-2.5*$10^6$ CTV-labelled (Invitrogen, C34557) naïve $CD8^+$ T cells were injected into the tail vein. For IL-7 blocking experiments, 0.5 µg anti-IL-7Ra antibody (A7R34, Bio X Cell, #BE0065) or isotype control (Bio X Cell, #BE0089) was added to the cell suspension and injected simultaneously in 100µl final volume. In Rag1KO and immunocompetent hosts, cells were left to expand for 5 and 14 days after transfer correspondingly.

Bone marrow chimeras

Mice were irradiated with 500cGy one day before the transplantation. The next day, the bone marrow was flushed out of the mouse hindlimb bones using a syringe filled with FACS buffer. For T cell depletion, bone marrow cells were suspended at a concentration of $10^7$ /ml, and hybridoma supernatants containing depletion antibodies (anti-CD4 – RL-172, anti-CD8 – 31M, anti-CD90 – T24) were added at volume ratios 1:10 to cell suspension. Cells were incubated on ice for 15 min, spun down (4 min 370g 4°), resuspended in Low Tox -M Rabbit complement

(Cedarlane, CL3051) at a concentration of $10^7$ /ml and incubated at 37°C for 45 min, washed with, spun down and resuspended in FACS buffer at a concentration 2.5 x $10^7$. After filtering the suspension, 5 x $10^6$ T cell-depleted bone marrow cells were injected into the tail vein. Engraftment in blood was monitored every 4 weeks until the final analysis of bone marrow, spleen, and blood 12 weeks post-transplantation.

EAE

EAE was performed in 6-8 weeks-old female mice. We induced EAE with MOG35-55 solution (Anaspec, AS-60130-10), mixed in a 1:1 ratio with CFA emulsion, containing M. tuberculosis (Difco, 231141) and IFA (Incomplete Freud's Adjuvant, BD or Sigma, 263910), with the final MOG35-55 peptide concentration of 1mg/ml.

Mice were injected 100 µl subcutaneously divided into both sides of the upper abdominal area, followed by intraperitoneal injection of 300 µl pertussis toxin solution, which was repeated on day 2 of the experiment. The severity score and weight were measured every 2 days for the first week of the experiment, then the severity score was measured daily. For mice reaching a score of 2.5, food was provided on the floor of the cage. On day 16, the spleen, axillary lymph nodes, and spinal cord were analyzed by flow cytometry.

Alfafold3

Amino acid sequences of M. musculus TMEM127 (Q8BGP5), SUSD6 (Q8BGE4), IL2RG (P34902), WWP2(Q9DBH0), without signal peptide, were processed using AlphaFold3 server with default parameters (*34*). The highest-ranking model was used for analysis and visualization using UCSF ChimeraX (*82*). For clarity and where appropriate, unstructured low-confidence regions were hidden from the field of view.

Statistical analysis

Was done using Prism 10 (GraphPad software) and R.

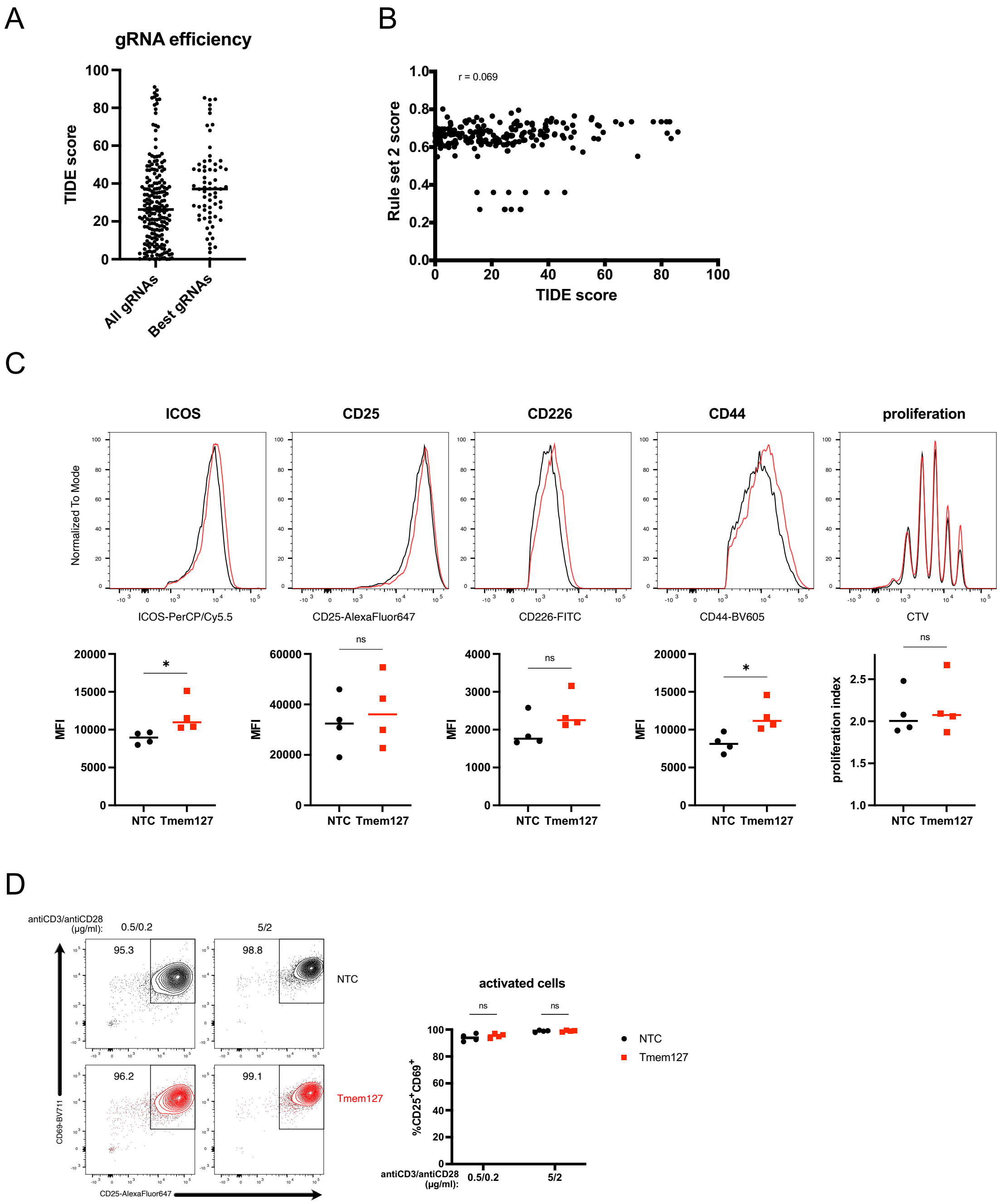
A
gRNA efficiency
TIDE score
All gRNAs
Best gRNAs
B
r = 0.069
Rule set 2 score
TIDE score
C
ICOS
CD25
CD226
CD44
proliferation
Normalized To Mode
ICOS-PerCP/Cy5.5
CD25-AlexaFluor647
CD226-FITC
CD44-BV605
CTV
MFI
proliferation index
NTC Tmem127
ns
*
D
antiCD3/antiCD28 (μg/ml):
0.5/0.2
5/2
95.3
98.8
96.2
99.1
NTC
Tmem127
CD69-BV711
CD25-AlexaFluor647
activated cells
%CD25+CD69+
NTC
Tmem127

**Fig. S1.** (**A**) Tracking of indels by Decomposition (TIDE) scores of gRNAs used in the CRISPR screening experiments, each dot represents single electroporation sample and line a median of one screening experiment, (**B**) Comparison of predicted (Rule set 2 score) with experimentally measured (TIDE) performance of gRNAs, (**C**) Activation markers and proliferation of 2 days in vitro activated naïve $CD4^+$ T cells after Tmem127KO or non-targeting control electroporation, (**D**) Frequency of activated Tmem127KO or non-targeting control naïve $CD4^+$ T cells after 2 days of in vitro stimulation with low (0.5/0.2 μg/ml) or high (5/2 μg/ml) concentration of anti-CD3/anti-CD28 antibodies, dots represent samples and line is a median of 4 (C, D) experiments. P values were calculated using an unpaired t-test or two-way ANOVA with Sidak's multiple comparison test. ns, not significant; *p < 0.05; **p < 0.01; ***p < 0.001; ****p<0.0001.

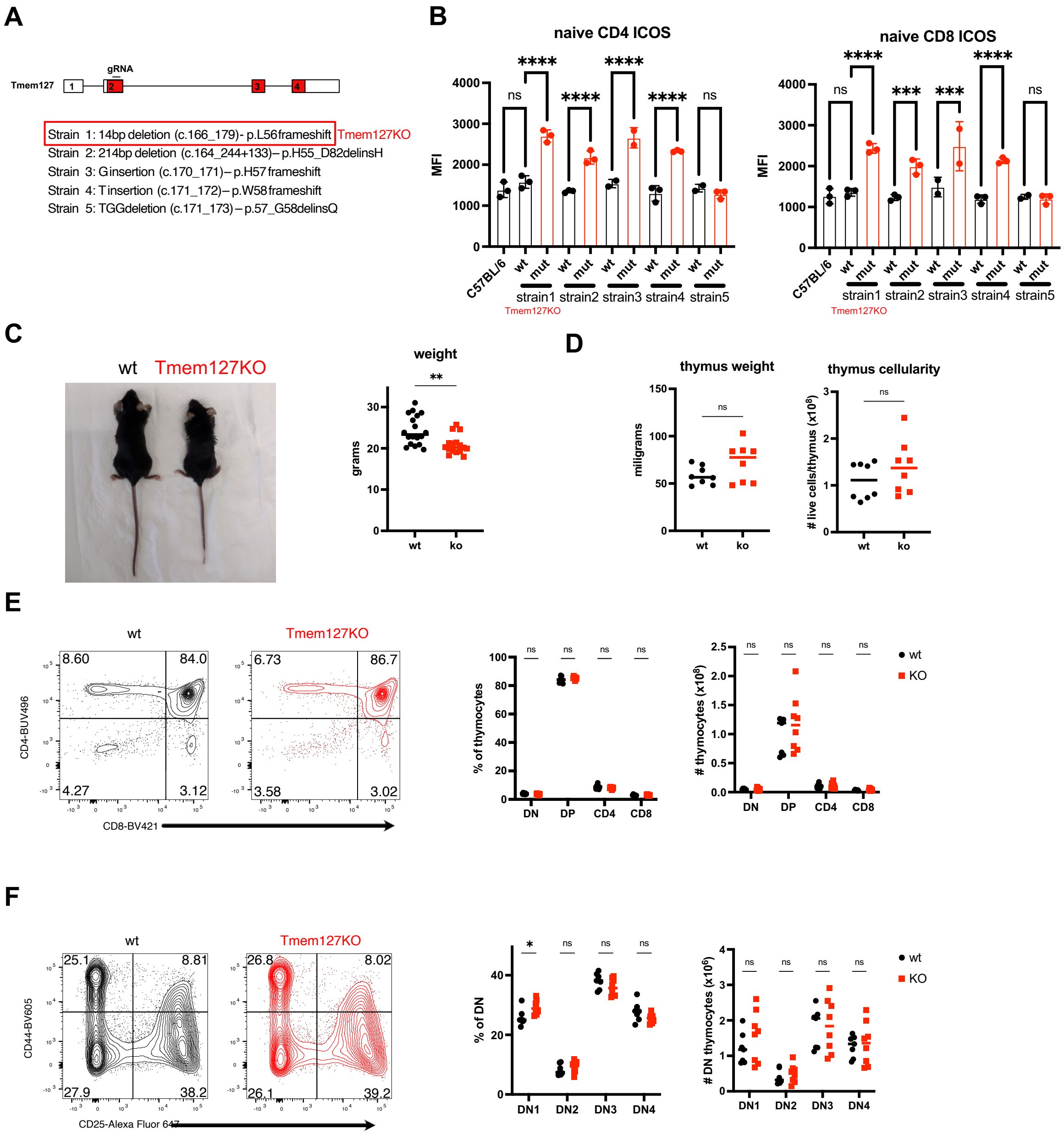
A
Tmem127
gRNA
Strain 1: 14bp deletion (c.166_179)- p.L56frameshift Tmem127KO
Strain 2: 214bp deletion (c.164_244+133)– p.H55_D82delinsH
Strain 3: G insertion (c.170_171)– p.H57 frameshift
Strain 4: T insertion (c.171_172)– p.W58 frameshift
Strain 5: TGG deletion (c.171_173) – p.57_G58delinsQ
B
naive CD4 ICOS
naive CD8 ICOS
MFI
C57BL/6
wt
mut
strain1
strain2
strain3
strain4
strain5
Tmem127KO
C
wt
Tmem127KO
weight
grams
ko
D
thymus weight
miligrams
thymus cellularity
### live cells/thymus (x10^8)
E
CD4-BUV496
CD8-BV421
% of thymocytes
### thymocytes (x10^8)
DN
DP
CD4
CD8
KO
F
CD44-BV605
CD25-Alexa Fluor 647
% of DN
### DN thymocytes (x10^6)
DN1
DN2
DN3
DN4

**Fig. S2.** (**A**) Summary of Tmem127 genome editing to create a germline knockout and generated mutant alleles, (**B**) Splenocytes from 5 mutant strains, their wt littermates and C57BL/6 controls were stained for ICOS, error bars represent mean and SD, (**C**) Photo representing size and quantification of body weight comparing Tmem127KO and wt mice, (**D**) Quantification of weight and cellularity of thymi isolated from Tmem127KO or wt mice, (**E-F**) Flow cytometry data and quantification of thymocyte populations isolated from Tmem127KO and wt mice and stained for CD4, CD8, CD25 and CD44, dots represent mice and lines median of 4 (B, D, E, F) or 5 (C) experiments. P values were calculated using unpaired t-test, one-way or two-way ANOVA with Sidak's multiple comparison test. ns, not significant; *p < 0.05; **p < 0.01; ***p < 0.001; ****p<0.0001.

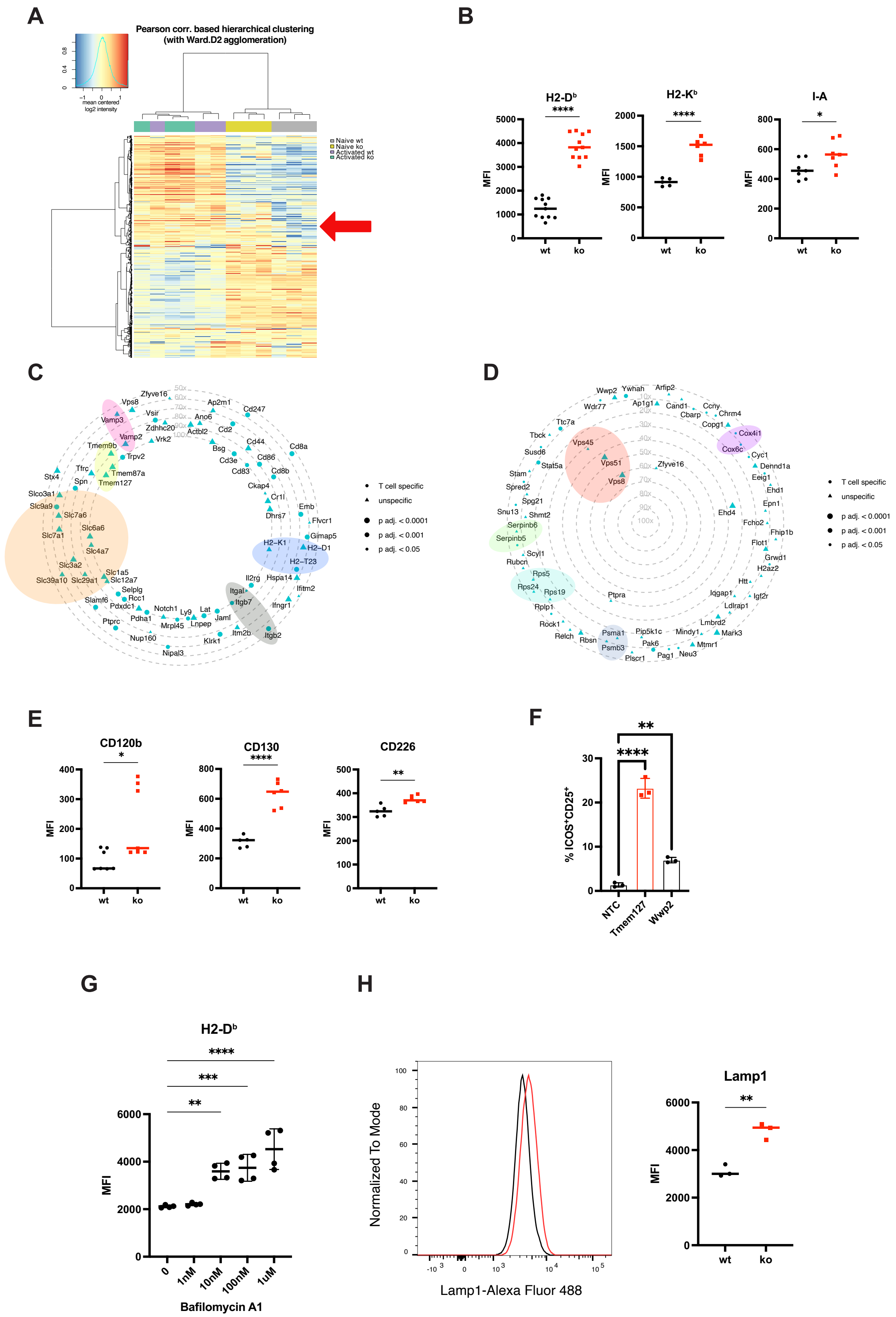
A
Pearson corr. based hierarchical clustering (with Ward.D2 agglomeration)
mean centered log2 intensity
Naive wt
Naive ko
Activated wt
Activated ko
B
H2-Dᵇ
H2-Kᵇ
I-A
MFI
wt
ko
C
D
T cell specific
unspecific
p adj. < 0.0001
p adj. < 0.001
p adj. < 0.05
E
CD120b
CD130
CD226
F
% ICOS⁺CD25⁺
NTC
Tmem127
Wwp2
G
H2-Dᵇ
Bafilomycin A1
0
1nM
10nM
100nM
1uM
H
Normalized To Mode
Lamp1-Alexa Fluor 488
Lamp1

**Fig. S3.** (**A**) Heatmap representing Pearson correlation based hierarchical clustering of proteomics data comparing naïve and 3 day in vitro activated naïve $CD8^+$ T cells isolated from Tmem127KO and wt mice, (**B**) Mean fluorescence intensities of indicated proteins stained in naïve $CD8^+$ T cells freshly isolated from Tmem127KO or wt mice, (**C**) Radar plot showing Bio-ID abundance ratios comparing detection in cells expressing full-length Tmem127-BirA versus the eGFP-BirA negative control, (**D**) Radar plot showing Bio-ID abundance ratios comparing detection in cells expressing full-length Tmem127-BirA versus the C-terminal truncated Tmem127-BirA negative control, (**E**) Mean fluorescence intensities of indicated proteins stained in naïve $CD8^+$ T cells freshly isolated from Tmem127KO or wt mice, (**F**) Frequency of $ICOS^+CD25^+$ naïve $CD4^+$ T cells electroporated with NTC, Tmem127- or Wwp2-targeting CRISPR RNPs and cultured for 5 days in medium containing 5 ng/ml IL-7, dots represent independent experiments, error bars represent mean of 3 experiments and SD, (**G**) Median fluorescence intensity of H2-$D^b$ staining on naïve $CD8^+$ T cells cultured in vitro for one day in presence of indicated concentrations of Bafilomycin A1, dots represent mice, line and error bars represent mean of 2 experiments and SD, (**H**) Flow cytometry data and quantification of intracellular Lamp1 levels in naïve $CD8^+$ T cells cultured in vitro for 5 days with 5 ng/ml IL-7, dots represent mice and lines median of 2 experiments. P values were calculated using an unpaired t-test or one-way ANOVA with Dunnett's multiple comparison test. ns, not significant; $*p < 0.05$; $**p < 0.01$; $***p < 0.001$; $****p<0.0001$.

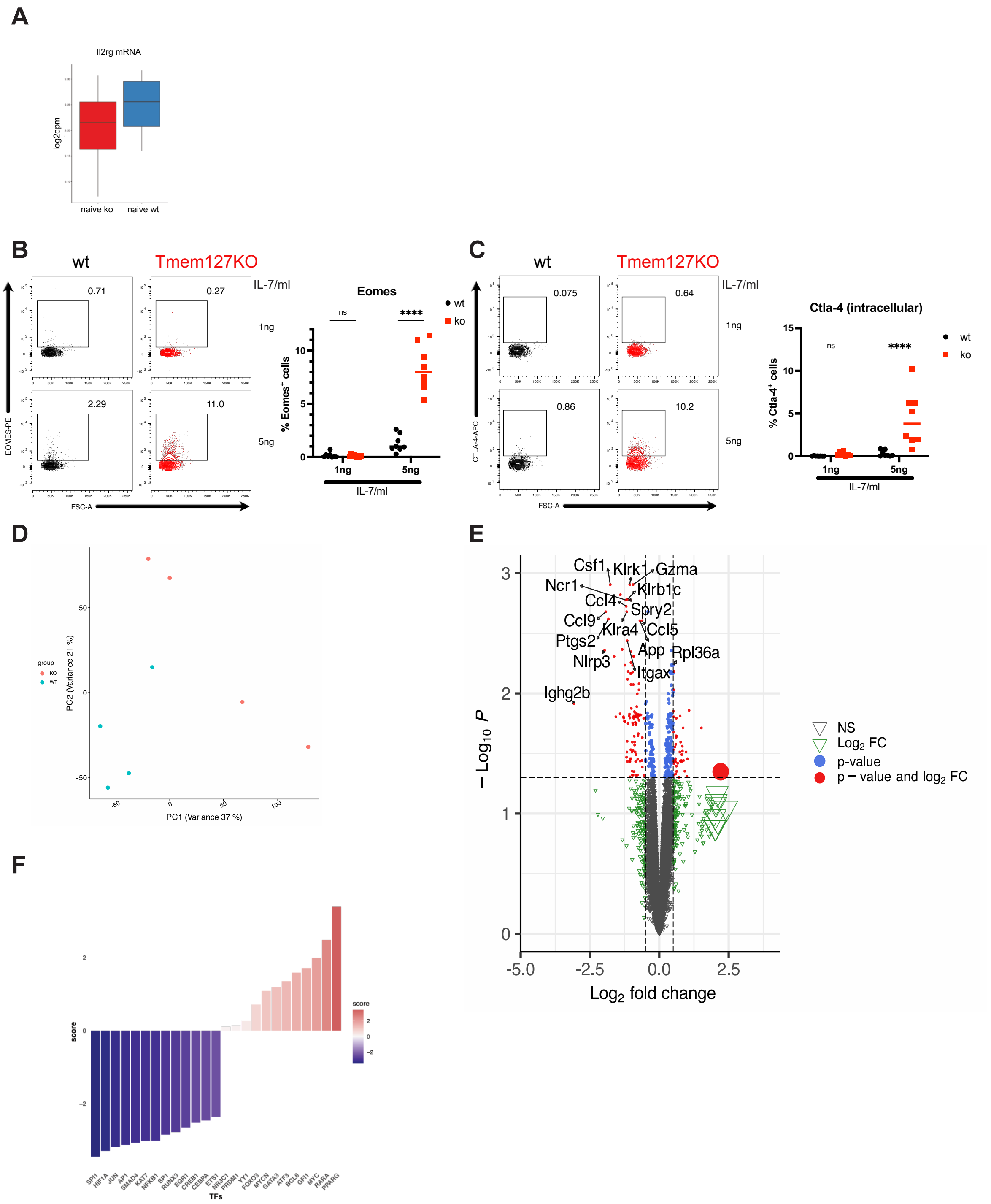
A
Il2rg mRNA
log2cpm
naive ko
naive wt
B
wt
Tmem127KO
IL-7/ml
0.71
0.27
1ng
2.29
11.0
5ng
EOMES-PE
FSC-A
Eomes
ns
****
wt
ko
% Eomes+ cells
1ng
5ng
IL-7/ml
C
wt
Tmem127KO
IL-7/ml
0.075
0.64
1ng
0.86
10.2
5ng
CTLA-4-APC
FSC-A
Ctla-4 (intracellular)
ns
****
wt
ko
% Ctla-4+ cells
1ng
5ng
IL-7/ml
D
group
KO
WT
PC2 (Variance 21 %)
PC1 (Variance 37 %)
E
Csf1
Klrk1
Gzma
Ncr1
Klrb1c
Ccl4
Spry2
Ccl9
Klra4
Ccl5
Ptgs2
App
Rpl36a
Nlrp3
Itgax
Ighg2b
− Log10 P
Log2 fold change
NS
Log2 FC
p-value
p − value and log2 FC
F
score
TFs
SPI1
HIF1A
JUN
AP1
SMAD4
KAT7
NFKB1
SP1
RUNX3
EGR1
CREB1
CEBPA
ETS1
NR3C1
PRDM1
YY1
FOXO3
MYCN
GATA3
ATF3
BCL6
GFI1
MYC
RARA
PPARG

**Fig. S4.** (**A**) RNAseq data of Il2rg mRNA expression in naïve $CD8^+$ T cells isolated from Tmem127KO and wt mice, (**B, C**) Intracellular levels and quantification of Eomes (B) and Ctla-4 (C) in naïve $CD8^+$ T cells isolated from Tmem127KO and wt mice and cultured in vitro for 4 days with 5 ng/ml IL-7, (**D**) Principal component analysis of RNAseq of naïve $CD8^+$ T cells isolated from Tmem127KO and wt mice, (**E**) Volcano plot showing differentially expressed genes between Tmem127KO and wt naïve $CD8^+$ T cells, (**F**) Transcription factors regulating differentially expressed genes between Tmem127KO and wt naïve $CD8^+$ T cells. Dots represent mice and line median of 2 experiments. P values were calculated using an unpaired t-test or two-way ANOVA with Sidak's multiple comparison test. ns, not significant; $^*p < 0.05$; $^{**}p < 0.01$; $^{***}p < 0.001$; $^{****}p<0.0001$.

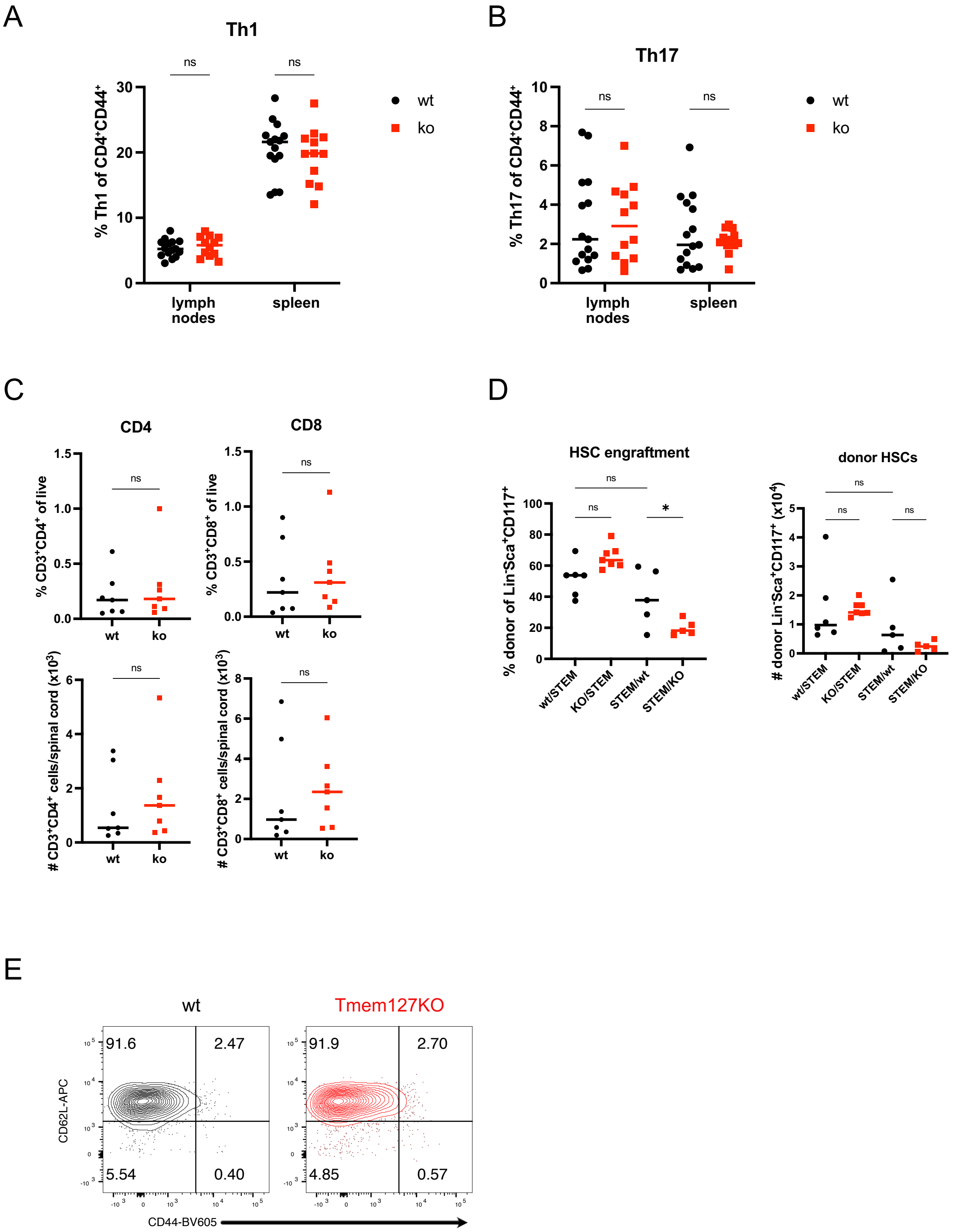
A
Th1
% Th1 of CD4+CD44+
ns
ns
lymph nodes
spleen
wt
ko
B
Th17
% Th17 of CD4+CD44+
ns
ns
lymph nodes
spleen
wt
ko
C
CD4
CD8
% CD3+CD4+ of live
% CD3+CD8+ of live
### CD3+CD4+ cells/spinal cord (x10³)
### CD3+CD8+ cells/spinal cord (x10³)
ns
wt
ko
D
HSC engraftment
donor HSCs
% donor of Lin-Sca+CD117+
### donor Lin-Sca+CD117+ (x10⁴)
ns
*
wt/STEM
KO/STEM
STEM/wt
STEM/KO
E
wt
Tmem127KO
91.6
2.47
5.54
0.40
91.9
2.70
4.85
0.57
CD62L-APC
CD44-BV605

**Fig. S5.** (**A**) Frequencies of CD44$^+$Tbet$^+$IFN$\gamma^+$ CD4$^+$ T cells in lymph nodes and spleens of Tmem127KO or wt mice at day 16 of $MOG_{35-55}$-induced EAE model, (**B**) Frequencies of CD44$^+$Ror$\gamma$t$^+$IL-17$^+$ CD4$^+$ T cells in lymph nodes and spleens of Tmem127KO or wt mice at day 16 of $MOG_{35-55}$-induced EAE model, (**C**) Frequencies and cell numbers of CD4$^+$ and CD8$^+$ T cells isolated from spinal cords of Tmem127KO or wt mice at day 16 of $MOG_{35-55}$-induced EAE model, (**D**) Frequencies of donor-derived neutrophils and dendritic cells in spleens of sublethally irradiated bone marrow chimeras, (**E**) Representative flow cytometry staining of donor naïve CD8$^+$ T cells isolated from spleens of recipient CD45.1 STEM mice 5 days after adoptive transfer. Dots represent mice and lines median of 2 experiments. P values were calculated using an unpaired t-test or two-way ANOVA with Sidak's multiple comparison test. ns, not significant; *$p < 0.05$; **$p < 0.01$; ***$p < 0.001$; ****$p<0.0001$.

#### References and Notes


67. J. Cuklina *et al.*, Diagnostics and correction of batch effects in large-scale proteomic studies: a tutorial. *Mol Syst Biol* **17**, e10240 (2021).
68. W. Stacklies, H. Redestig, M. Scholz, D. Walther, J. Selbig, pcaMethods--a bioconductor package providing PCA methods for incomplete data. *Bioinformatics* **23**, 1164-1167 (2007).
69. A. Dobin *et al.*, STAR: ultrafast universal RNA-seq aligner. *Bioinformatics* **29**, 15-21 (2013).
70. Y. Liao, G. K. Smyth, W. Shi, featureCounts: an efficient general purpose program for assigning sequence reads to genomic features. *Bioinformatics* **30**, 923-930 (2014).
71. M. D. Robinson, D. J. McCarthy, G. K. Smyth, edgeR: a Bioconductor package for differential expression analysis of digital gene expression data. *Bioinformatics* **26**, 139-140 (2010).
72. M. D. Robinson, A. Oshlack, A scaling normalization method for differential expression analysis of RNA-seq data. *Genome Biol* **11**, R25 (2010).
73. I. M. P. Badia *et al.*, decoupleR: ensemble of computational methods to infer biological activities from omics data. *Bioinform Adv* **2**, vbac016 (2022).
74. D. Turei, T. Korcsmaros, J. Saez-Rodriguez, OmniPath: guidelines and gateway for literature-curated signaling pathway resources. *Nat Methods* **13**, 966-967 (2016).
75. C. Schultheiss *et al.*, A20 haploinsufficiency disturbs immune homeostasis and drives the transformation of lymphocytes with permissive antigen receptors. *Sci Adv* **10**, eadl3975 (2024).
76. D. A. Bolotin *et al.*, MiXCR: software for comprehensive adaptive immunity profiling. *Nat Methods* **12**, 380-381 (2015).
77. M. Choi *et al.*, MSstats: an R package for statistical analysis of quantitative mass spectrometry-based proteomic experiments. *Bioinformatics* **30**, 2524-2526 (2014).
78. O. Makarova, E. Kamberov, B. Margolis, Generation of deletion and point mutations with one primer in a single cloning step. *Biotechniques* **29**, 970-972 (2000).
79. J. R. Wisniewski, A. Zougman, N. Nagaraj, M. Mann, Universal sample preparation method for proteome analysis. *Nat Methods* **6**, 359-362 (2009).
80. A. Grosche *et al.*, The Proteome of Native Adult Muller Glial Cells From Murine Retina. *Mol Cell Proteomics* **15**, 462-480 (2016).
81. L. Molitor *et al.*, Depletion of the RNA-binding protein PURA triggers changes in posttranscriptional gene regulation and loss of P-bodies. *Nucleic Acids Res* **51**, 1297-1316 (2023).
82. T. D. Goddard *et al.*, UCSF ChimeraX: Meeting modern challenges in visualization and analysis. *Protein Sci* **27**, 14-25 (2018).
